# mutscan - a flexible R package for efficient end-to-end analysis of multiplexed assays of variant effect data

**DOI:** 10.1101/2022.10.25.513674

**Authors:** Charlotte Soneson, Alexandra M Bendel, Guillaume Diss, Michael B Stadler

## Abstract

Multiplexed assays of variant effect (MAVE) experimentally measure the fitness of large numbers of sequence variants by selective enrichment of sequences with desirable properties followed by quantification by sequencing. *mutscan* is an R package for flexible analysis of such experiments, covering the entire workflow from raw reads up to statistical analysis and visualization. Core components are implemented in C++ for efficiency. Various experimental designs are supported, including single or paired reads with optional unique molecular identifiers. To find variants with changed relative abundance, *mutscan* employs established statistical models provided in the *edgeR* and *limma* packages. *mutscan* is available from https://github.com/fmicompbio/mutscan.

## Background

A major question in biology is that of how sequence and function are related. The advances made in modern sequencing technology have resulted in an exponential increase in whole-genome and exome sequencing data over the past few decades and Genome Wide Association Studies (GWAS) have found statistical associations between certain genetic variants and phenotypes or diseases (1). However, the phenotypic consequences of a large fraction of variants identified in the human genome remain elusive (2), which is why these variants have been termed variants of uncertain significance (VUS). For example, 41.8 % of variants currently listed in ClinVar are characterized as VUS (3). Therefore, a pressing objective has been to find ways to annotate these variants in an efficient way.

Over the past decades, Multiplexed assays of variant effect (MAVE) have revolutionized the study of sequence-function relationships by enabling the simultaneous assessment of the functional consequences of thousands of sequence variants on a given phenotype. For example, a large library of variants is created by mutating a sequence of interest (deep mutational scanning, DMS), and this library is exposed to a pooled selective assay which results in an enrichment of variants with high activity in the given assay and a depletion of variants with low activity (4). The frequency of each variant before and after selection can be quantified using high throughput sequencing (Figure 1A). Variant counts can be obtained by either sequencing the variants directly or using molecular barcodes that uniquely identify each variant. The latter can reduce sequencing cost and increase read quality (5). Enrichment scores calculated from the variant frequencies can be used to infer molecular function and thus the functional effect of a mutation relative to the wildtype sequence (4). The variety of experimental designs that can be used in MAVE emphasize the value of these assays and their flexibility in addressing diverse biological questions. They have been used to examine activities of proteins, such as protein-protein interaction (PPI) (6–8), E3 ubiquitin ligase activity (8), protein abundance (7,9), receptor binding (10), aggregation (11, 12), and activity within signaling pathways (13). The functional assays used to achieve enrichment or depletion of variants are equally diverse and include for example fitness or cell growth (6,7,11,12), different reporter assays coupled with fluorescence-activated cell sorting (FACS) (9,10,13,14), and protein display (8, 15).

**Figure 1.**
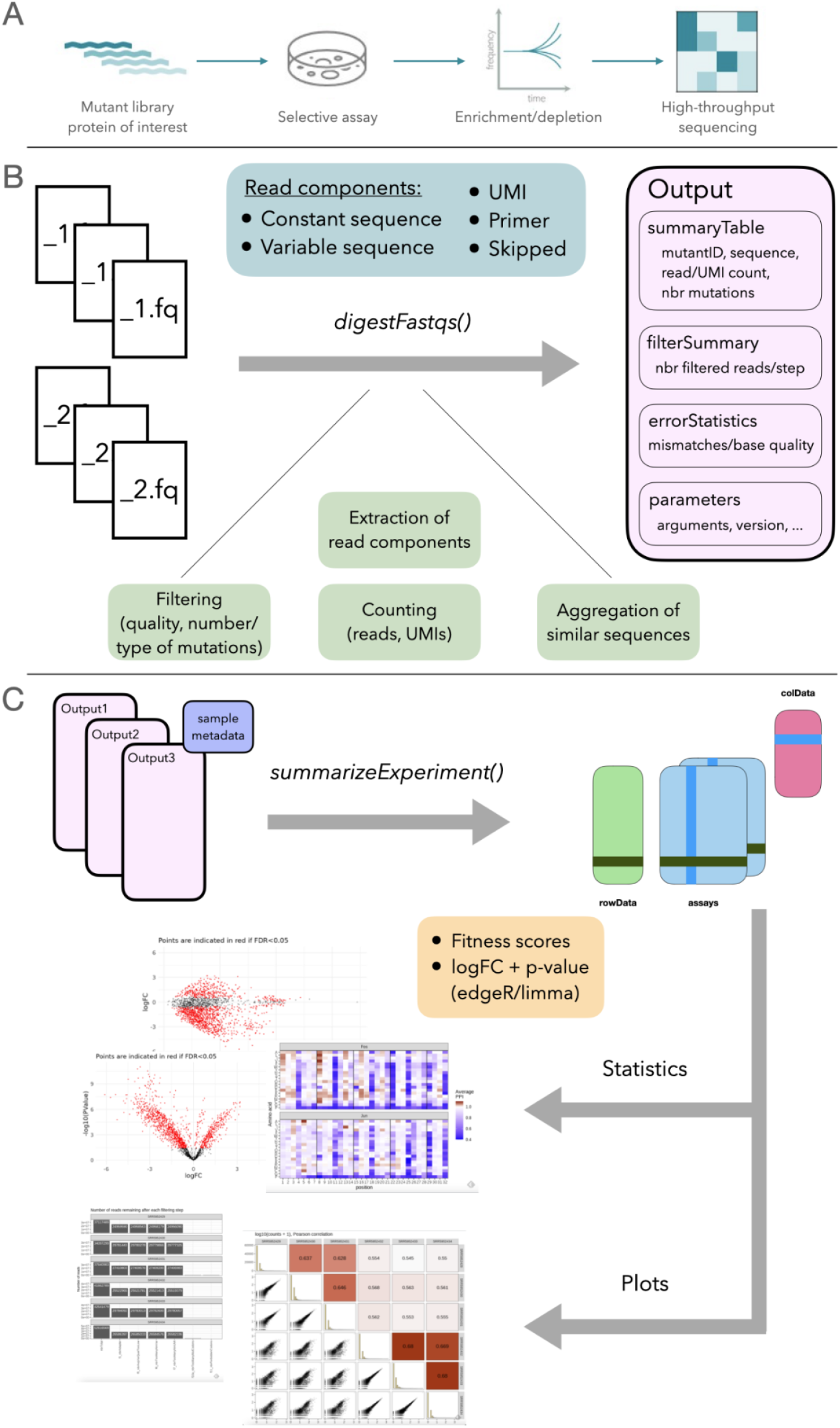
Overview of the main functionality of *mutscan*. **A**. Multiplexed Assays of Variant Effect experiments are based on the enrichment and depletion of protein variants in an assay that selects for a desired activity of the given variants. Enrichment and depletion are quantified using high-throughput sequencing. **B**. The *digestFastqs()* function processes the FASTQ file(s) for each sample independently. **C**. The output objects are then combined into a joint *SummarizedExperiment* object which is in turn provided to downstream functions for statistical analysis and plotting. For more details about the individual steps, see the main text.

The growth of the field has been further driven by the decreasing cost of sequencing and the simplified construction of large libraries thanks to commercial large-scale synthesis of DNA oligonucleotides (16). New technical developments that allow the synthesis of large libraries of entire synthetic genes will probably result in even larger libraries (17). Recently, a database was launched with the aim to collect the rich data gathered from MAVE assays in a central place with a unified structure to make it accessible for the scientific community (18). This, however, also calls for streamlined and more standardized analysis methods, including rigorous statistical analysis that considers the possible sources of error in MAVE experiments and therefore allows to make confident statements about the true functional consequences of variants. Several tools have been published to address this demand (Table 1), the most elaborate and widely used amongst these are *Enrich2* (19) and *DiMSum* (20).

**Table 1.** Comparison of different tools for the analysis of MAVE data. In our evaluations, we compare *mutscan* to *DiMSum* and *Enrich2*, as these are widely used in the field, and align with *mutscan* in terms of their scope and aim.

| Aspect | *mutscan* | *DiMSum* (20) | *Enrich2* (19) |
| --- | --- | --- | --- |
| OS availability | All | Unix-like | All (requires Python 2.7, end of life since Jan 1, 2020) |
| External dependencies (in addition to R/python packages) | None | FastQC (21), cutadapt (22), VSEARCH (23), starcode (24) | None |
| Diagnostic report | Yes | Yes | Yes (autogenerated plots) |
| Parallelizable | Yes | Yes | No |
| Supported library design | Single-end, paired-end | Single-end, paired-end | Single-end, paired-end |
| Statistical test specification | Any fixed effect model (formula) | No test. Calculates a fitness score and error, limited to 9 replicates | One score per replicate, combined using a mixed-effects model |
| Modularity | Samples are processed independently. Modularity between workflow steps. | The entire experiment is processed together. Modularity between workflow steps if intermediate files are retained. | The entire experiment is processed together. Intermediate results are reused if present. |
| Installation package | R package | Bioconda package | None |
| Enrichment scores | Absolute and relative | Relative | Absolute and relative |
| Use of barcodes or variant sequence | Both | Both | Both |
| Alignment strategy | Tree-based string matching (Hamming or Levenshtein distance) | Clustering (Levenshtein distance, using starcode (24)) | String matching or gapped alignment (Needleman-Wunsch) |
| UMIs | Yes | No | No |
| Filtering options | Read length consistency, forbidden codons, base quality, nbr mismatches in constant sequence, nbr mutated bases/codons, nbr N bases | Minimal read length, base quality, expected errors, nbr mutated bases/amino acids, min nbr reads, invalid overlap, forbidden substitution, mutation type | Base quality, min nbr reads, nbr mutations, nbr N bases, invalid overlap |
| Multiple amplicons | Yes | No | No |

Here, we present *mutscan*, a novel R package that provides a unified, flexible interface to the analysis of MAVE experiments, covering the entire workflow from FASTQ files to count tables and statistical analysis and visualization. The core read processing module is implemented in C++, which enables the analysis of large sequencing experiments within reasonable time and memory constraints. *mutscan* is easy to install and use, has a flexible interface that encompasses a broad range of experimental designs, and employs established statistical testing frameworks developed for count data. We apply *mutscan*, as well as *Enrich2* and *DiMSum*, to several experimental MAVE data sets and show that while estimated counts and fitness scores are often highly concordant between methods, *mutscan* is generally able to process the data faster, with lower memory requirements, and more efficient use of multi-core processing. Given the variety of MAVE experimental designs, the ever increasing scale of MAVE experiments and the democratization of the field, we believe that its flexibility, efficiency and ease of access will make *mutscan* an important addition to the MAVE analysis tool ecosystem.

## Results and discussion

### Example data sets

The results presented below are obtained by applying *mutscan* and other tools (Table 1) to four deep mutational scanning data sets (Table 2). These data sets represent a variety of typical MAVE experimental designs, and have been previously used for evaluation purposes (20).

**Table 2:**
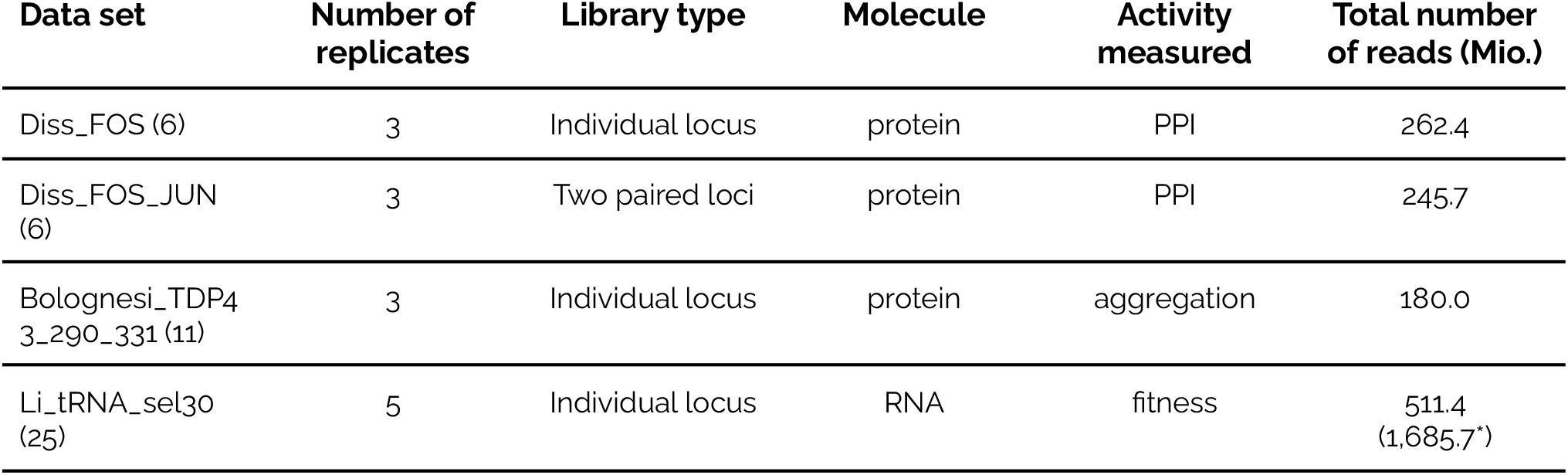
Overview of deep mutational scanning data sets used in this study. *In the Li_tRNA_sel30 data set, a single input replicate is paired with multiple output replicates. While the modular design of *mutscan* allows to process this shared input replicate just once, *DiMSum* (with the design used here, in agreement with (20)) requires it to be processed repeatedly, once for each selected replicate, which increases the number of processed read pairs from 511.4 to 1,685.7 Mio.

### Overview of the *mutscan* workflow

*mutscan* is implemented as an R package, with core processing modules written in C++ for efficiency. Figure 1B-C provides an overview of the full analysis workflow (for more details about the individual steps, see Methods). The first part of the workflow processes each sequencing library independently. Hence, additional samples can easily be added to an experiment without the need to re-process the existing samples. In this step, reads that do not adhere to the user specifications (e.g., too low base quality, too many or forbidden mutations compared to a provided wildtype sequence) are filtered out, and the remaining ones are used to tabulate the number of reads (or unique UMI sequences, if applicable) corresponding to each observed sequence variant. For increased processing speed, this step can be parallelized. In the second part, the output from all samples in the experiment are combined into a joint *SummarizedExperiment* object (26), containing the merged count matrix, a summary of the filtering applied to each individual sample, and additional information about the detected variants, such as the nucleotide and amino acid sequence, the number of mutated bases, codons and amino acids, and the type of mutations (silent, non-synonymous, stop). Finally, the merged object can be used as the input to functions generating diagnostic plots and reports, as well as statistical analysis functions that estimate fitness scores and find variants that are increasing or decreasing significantly in abundance during the selection process. Since the data is represented as a *SummarizedExperiment* object, it can also be directly used as input to a wide range of analysis and visualization tools from the Bioconductor ecosystem (27).

### Case study: evaluating interaction strength between FOS and JUN variants

To illustrate the practical use of *mutscan*, we reprocess the 'trans' data set from (6). Here, libraries of single amino acid mutants of FOS and JUN’s basic leucine zipper domains were constructed by oligonucleotide replacing each codon by one of the 32 codons ending with a C or a G (encoded by NNS in the IUPAC code (28)). The two libraries were then cloned together on the same plasmid to measure the effects of combining one mutation on each partner on the protein-protein interaction (PPI) between FOS and JUN. Interaction was scored by deepPCA (6), which couples protein-protein interaction to growth rate in a typical MAVE setting. A rendered report detailing the full analysis can be found in Additional file 1. The data set contains three replicates, each with an input and an output sequencing library (before/after the selection assay, respectively). As indicated above, we start by processing each of the six FASTQ files separately with the *digestFastqs()* function from *mutscan*. This step extracts the sequences of the variable regions corresponding to the FOS and JUN variants from the paired reads, compares them to the provided wildtype sequences and identifies the differences, and tabulates the number of reads and unique UMIs for each identified variant combination. Since the variable regions of the forward and reverse reads correspond to variants of different proteins encoded at two different loci and do not share any common sequence, we instruct *digestFastqs()* to process them separately rather than attempting to merge them. This also allows us to submit separate wildtype sequences for the variable regions in the forward and reverse reads. We retain only reads with at most one mutated codon in each of the two proteins. The output of this initial processing step is a list for each sample, containing a count table, a filtering summary and a record of the parameters that were used (Fig. 1B). While these objects can be explored as they are, it is more convenient to merge them into a joint *SummarizedExperiment* (26) object for downstream analysis (Fig. 1C), which is done via the *summarizeExperiment()* function in *mutscan*. The resulting object contains a matrix with the UMI counts for all variants in all six samples, as well as a summary of the number of reads filtered out at each step, and any metadata provided for the samples (replicate ID, condition, optical density, etc), and feeds directly into the diagnostic plot and statistical analysis functions in *mutscan*, including *plotPairs()* and *plotFiltering()*, the outputs of which are shown in Figure 2. We observe that, as expected, we find more unique variants with multiple base mutations, but the observed abundance of individual variants with multiple mutations is markedly lower than for variants with no or a single mutated base (Figure 2A-B). Using *mutscan* to visualize the filtering process further illustrates that across all samples, the main reasons for read pairs being filtered out are that they contain an adapter, or that they contain more than the allowed number of mutated codons (Figure 2C-D).

**Figure 2.**
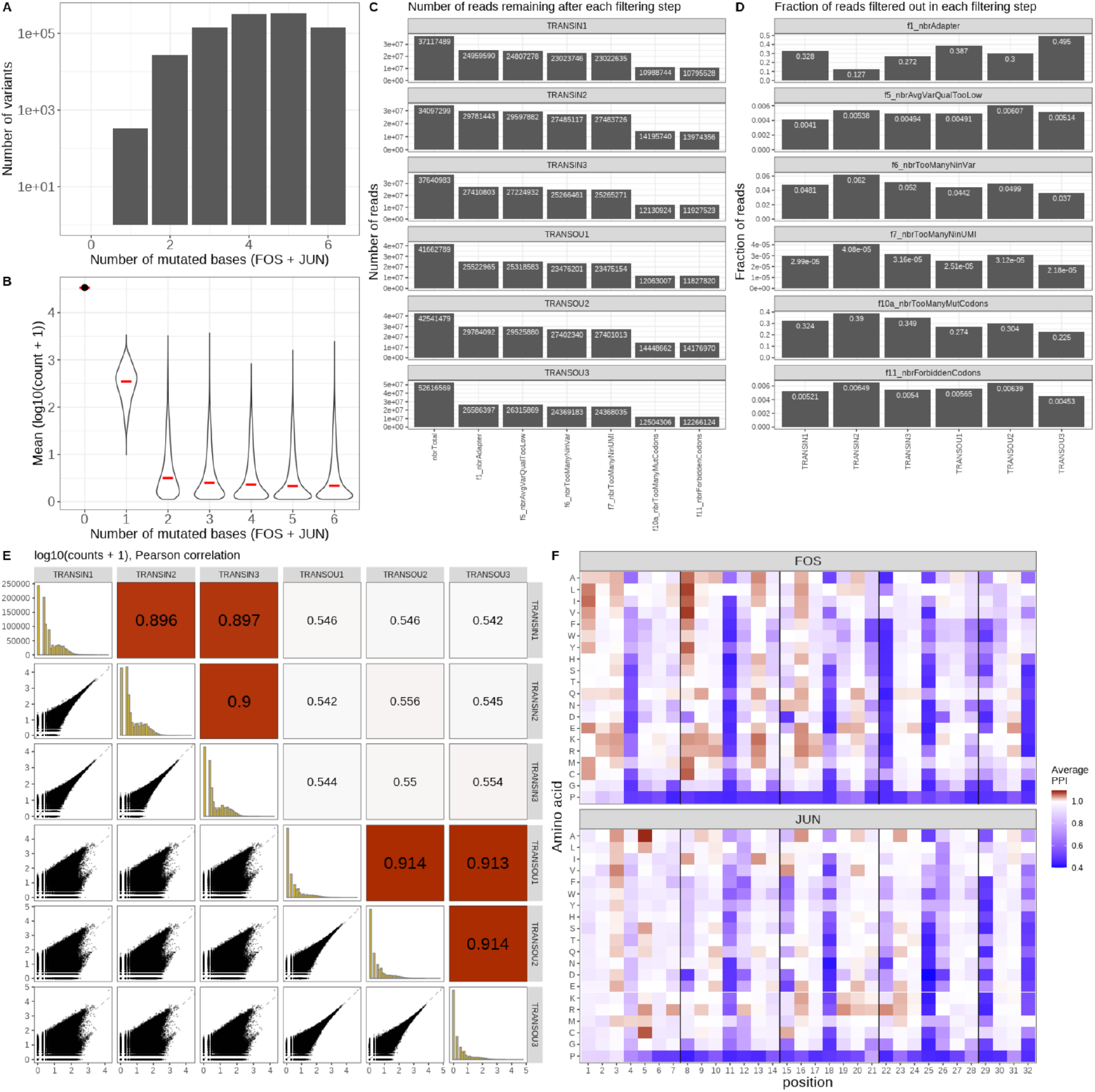
Results from the *mutscan* re-analysis of the FOS/JUN protein interaction data set from (6). **A**. The number of variants detected with different numbers of mutated bases. **B**. The average abundance of variants with different numbers of mutated bases. **C-D**. Diagnostic plots of read filtering performed by *digestFastqs()*. **E**. Pairs plot displaying the concordance of the observed counts across the six samples. **F**. Heatmap showing the estimated protein-protein interaction score for all single-amino acid variants of each of the two proteins.

Next, we use *mutscan* to investigate the concordance among the six samples, by plotting the estimated variant counts (Figure 2E). As expected, the correlation within each type of sample (input/output) is considerably higher than the correlation between input and output samples, indicating that the selection step indeed influences the sample composition.

After the initial quality assessment, we use *mutscan* to summarize the counts for variants with the same amino acid sequence. From this matrix, we then estimate a protein-protein interaction score for each variant and replicate, indicating the fitness of the variant relative to the wildtype sequence as described by (6). Focusing only on variants with a mutated amino acid in either FOS or JUN (but not both), we can generate a heatmap summarizing the impact of each single amino acid mutation on the overall interaction score (Figure 2F).

### *mutscan* enables processing flexibility

The *digestFastq()* function in *mutscan*, which performs the initial processing of each individual library, is designed to provide flexibility in the sample processing, thereby enabling the analysis of a wide range of library designs, not limited to MAVE experiments. Here, we highlight some of the main features.

- *Processing of single- or paired-end reads with arbitrary composition of basic elements* : *mutscan* accepts FASTQ files from both single-end and paired-end experiments. In addition, a vector of (pairs of) FASTQ files can be provided and these will be internally concatenated. If some part of the variable regions of the forward and reverse reads is shared, e.g., if they correspond to overlapping parts of the same protein, the reads in a pair can be merged before further processing. The size of the overlap can be anywhere between a single base and the length of the whole read, and restrictions on what constitutes a valid overlap can be specified by the user. For each read, the user further specifies the sequence composition in terms of five element types (see Box 1): variable regions (typically the sequences of interest), constant regions, skipped regions, primers, and UMIs. Each read can contain multiple (adjacent or nonadjacent) regions of each element type except primers, and the lengths of the regions can be defined by the user or automatically inferred by *mutscan*. This design provides an intuitive interface for the user, and implies that many different types of experiments can be analysed within the same framework. Moreover, processing parameters and read compositions are specified separately for the forward and reverse reads, which allows direct processing of constructs with multiple variable regions, e.g., corresponding to different proteins.
- *Sequence-based analysis or comparison (or collapsing) to one or more wildtype sequences*: *mutscan* allows the optional specification of one or more wildtype sequences, against which the extracted variable regions will be compared. If more than one wildtype sequence is provided, *mutscan* will find the most similar one for each read, and match the read against that. The variant identifiers used by *mutscan* consist of the name of the most similar wildtype sequence, augmented with the observed deviation(s). For example, an identifier of the form GENEX.10.A indicates that the closest wildtype sequence was that of GENEX, and the observed read deviated from this wildtype sequence in that the tenth base was an A, rather than the reference base. It is also possible to collapse all variants to their closest reference sequence, if the mutations are not of interest. If no wildtype sequence is provided, *mutscan* represents each identified variant by its actual nucleotide sequence.
- *Codon- or nucleotide-based analysis*: *mutscan* allows processing of both coding and non-coding sequences. If wildtype sequences are provided, the user can choose to limit either the number of mutated bases or the number of distinct mutated codons that are allowed in the identified variants. This choice will further impact the naming of the variants (in terms of the codon or nucleotide deviations from the closest wildtype sequence).
- *Collapsing of similar sequences*: If no wildtype sequences are provided, the user has the option to collapse variants (unique variable sequences) with at most a given number of mutations between them. The collapsing is done in a greedy way, starting from the most abundant variant, and can be limited to only collapsing variants with a large enough abundance ratio.
- *Processing only a subset of the reads*: For testing purposes, it is often useful to process only a small subset of the reads. *mutscan* allows the user to limit the processing to the first N reads in the FASTQ file, where N is specified by the user.
- *Various filtering criteria*: *mutscan* implements a range of filtering criteria, including the number of 'N' bases in the variable and/or UMI sequence, the number of mutations in the constant and/or variable sequence (if a reference sequence is provided), the base quality of the identified mutations and/or the average base quality in the read, the presence of forbidden codons (specified using IUPAC code), or invalid overlap between forward and reverse reads for merging. The output object contains a table listing the number of reads filtered out by each of the criteria.
- *Export of excluded reads (with reason for exclusion) to FASTQ files for further investigation*: In cases where reads are filtered out for any of the reasons listed above, it may be helpful to be able to process these further. *mutscan* can write reads that are filtered out to a (pair of) FASTQ file(s), including the reason for exclusion in the read identifier.
- *Estimation of sequencing error rate*: If the input reads contain a constant region, *mutscan* estimates the sequencing error rate by counting the number of mismatches compared to the expected sequence across the reads. The error rate is further stratified by the base quality reported in the FASTQ file.
- *Nucleotide- or amino acid-based analysis*: The main output from *digestFastqs()* is represented in base or codon space. However, the corresponding amino acid information is recorded, and the count matrix can easily be collapsed on the amino acid level. In addition, *mutscan* reports the type of mutations (silent, non-synonymous, stop) present in each variant.
- *Flexible analysis frameworks*: The statistical testing module in *mutscan* is based on established packages for analysis of digital gene expression data (*edgeR* (29) and *limma-voom* (30)). Both tools allow the user to specify an arbitrary (fixed-effect) design, and thus provides excellent flexibility for testing complex hypotheses, not limited to paired comparisons of input and output samples. Moreover, several different normalizations are available, allowing calculation of both 'absolute' and 'relative' log-fold changes (e.g., changes relative to a wildtype reference).

##### Box 1. Read components

*mutscan* requires the user to specify the composition of the input read(s) in terms of the following five component types:

- variable regions (V): these are typically the regions of interest. If one or more wildtype sequences are provided, the variable regions will be compared to those to identify variants. If no wildtype sequences are provided, the sequence of the variable region will be used to represent the variant.
- constant regions (C): these regions are used to estimate sequencing error rates. Reads can also be filtered out if they have too many mutations in the constant region.
- skipped regions (S): these regions will be ignored in the processing.
- primers (P): a primer sequence differs from a constant region by the fact that it is not required to occur in a pre-defined position in the read. Instead, *mutscan* will search the read for a perfect match for the primer sequence.
- UMIs (U): these sequences will be used to correct for PCR amplification biases. If present, the output count table will contain, for each variant, both the number of reads and the number of unique associated UMIs.

For example, an experiment where the reads have the following composition:

**[1 skipped nt] - [10 nt UMI] - [18 nt constant sequence] - [96 nt variable region]**

would be specified to *digestFastqs()* by an element string "SUCV" and an element length vector c(1, 10, 18, 96).

For a library design with a primer sequence, the primer acts as an ‘anchor’, and the read composition before and after the primer is specified. For example, reads with the following composition:

**[unknown sequence] - [10 nt primer] - [variable region, constituting the remainder of the read]**

would be represented by an element string "SPV" and an element length vector c(−1, 10, −1), where the −1 indicates that the corresponding read part consists of the remaining part of the read, not accounted for by any of the other specified parts.

### *mutscan* controls the type I error

To evaluate whether the statistical approaches employed by *mutscan* are able to control the type I error rate at the expected level, we designed a null comparison for each of the example data sets. Briefly, we assigned approximately half of the replicates to group 'A', and the rest of the replicates to group 'B', and tested, for each variant, whether the output/input count ratio was different between the two artificial groups. The procedure was repeated for all possible assignments of the replicates to two approximately equally sized groups. Overall, the nominal p-value distributions from *mutscan*, using either *edgeR* and *limma* as the inference engine, were largely uniform, indicating that the tests are well calibrated (Figure 3).

**Figure 3.**
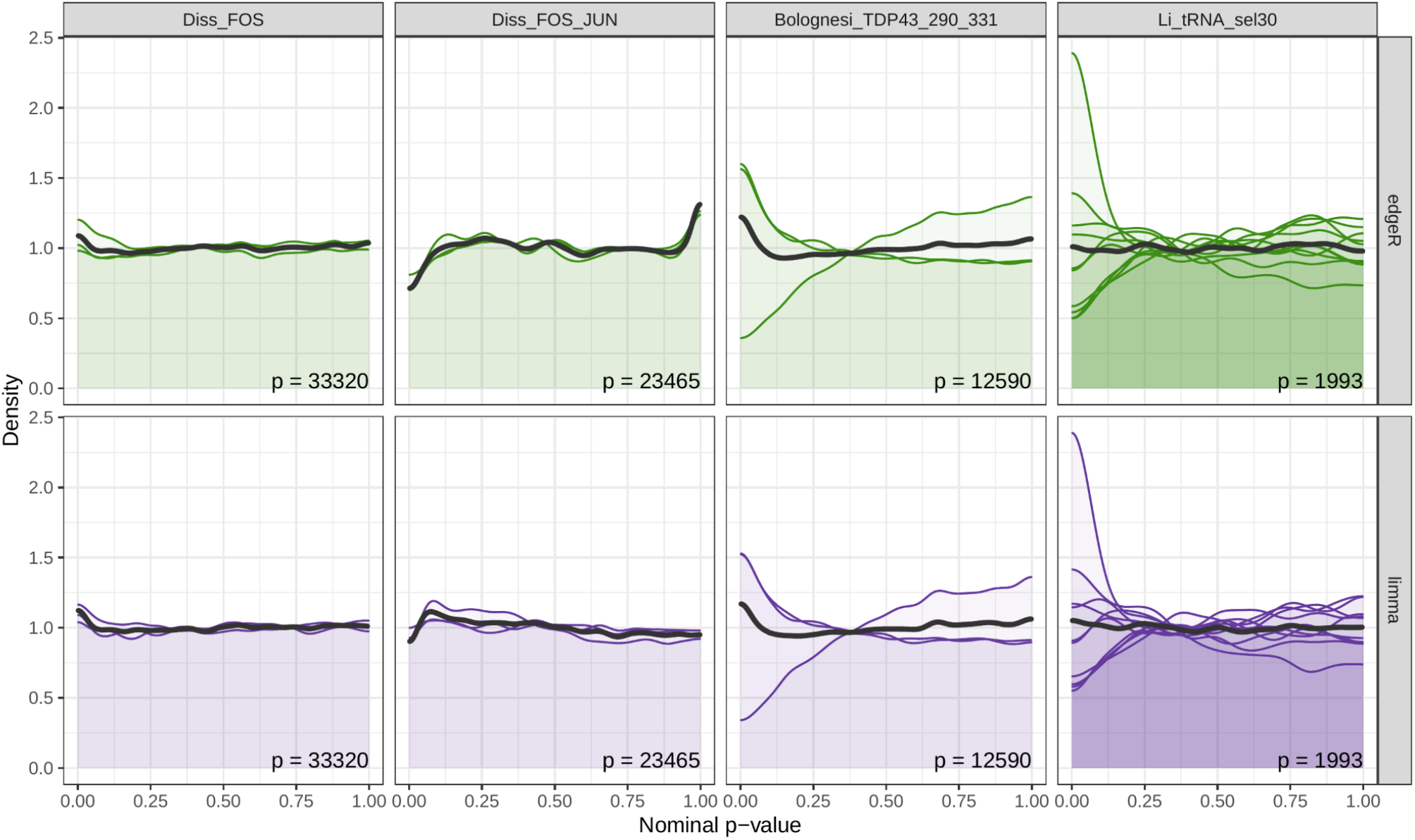
*mutscan* p-value distributions for null comparisons. For each data set, repeated null data sets were generated by artificially splitting the replicates into two approximately equally sized groups. For each such artificial null data set, *mutscan* (with method set to *edgeR* and *limma*, respectively) was used to fit a model and test whether the log-fold change between input and output samples differed significantly between the two artificial groups. The colored densities represent the individual data splits, while the dark grey density represents the union of p-values from all data splits. Since the groupings are artificial, uniform p-value distributions are expected. While technical differences among the samples, and the low sample size in general, imply that not all comparisons provide exactly uniform p-value distributions, we do not observe a systematic bias in the p-values from *mutscan*. Only variants with more than 50 counts in all input samples were considered for this analysis. The number of retained features is indicated in each panel.

### *mutscan* is computationally efficient

In order to evaluate the computational efficiency of *mutscan* (v0.2.31), we applied it to the four example data sets and used the benchmarking capabilities of *snakemake* (31) to track CPU and memory usage as well as the volume of data input and output. We also ran *DiMSum* v1.2.11 (on all data sets) and *Enrich2* v1.3.1 (for the Diss_FOS data set only, due to the long execution time), attempting to set the parameters of the different tools in such a way that the similarity between the performed analyses would be maximized (see Methods). We evaluated the entire process, from raw FASTQ files to output tables, and additionally included the generation of the summary report where applicable.

For all data sets, the total execution time for *mutscan* was lower than for *DiMSum* (Figure 4), and both of these finished in less time than *Enrich2*. The distribution of time spent in the different stages varied across data sets as the processing time for individual samples is largely determined by the number of reads, while the time required to perform statistical tests and generate plots for quality assessment is rather dependent on the number of identified and retained variants. The same effect is seen for the memory consumption. Only a small amount of memory is required for the initial sample processing by *mutscan*, while the memory required for the later analysis and statistical testing depends on the number of retained variants, and thus the size of the count matrix that needs to be loaded into the R session for processing. We also note that, apart from reading the FASTQ files, the total volume of input and output data for *mutscan* is very low, as the whole sample processing is performed by a single function that traverses the input files only once and without the need to create intermediate files on disk for transfer of data between different software tools. The total runtime of *DiMSum*, which is reading and writing much larger volumes of data to disk, also likely depends more strongly on the performance of the storage system. Another consequence of *mutscan*’s design is that a larger fraction of the time is spent on actions that are parallelizable, which can be seen by its higher average CPU load (400-600% for *mutscan* when run with 10 cores, to compare with 100-200% for *DiMSum* with 10 cores). *Enrich2* is not parallelizable and thus exhibits a constant load of 100%.

**Figure 4.**
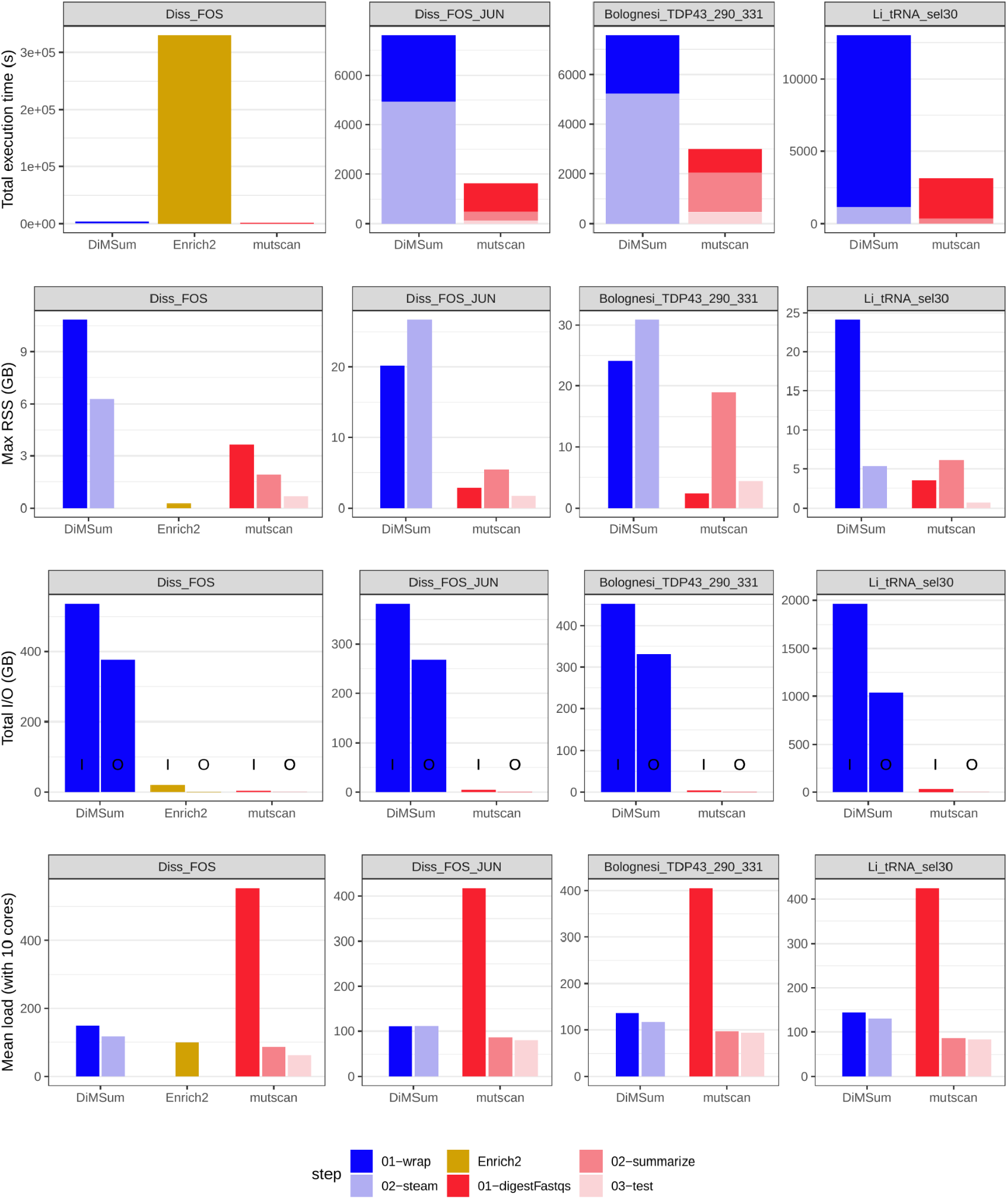
Comparison of computational performance metrics for *mutscan*, *Enrich2* and *DiMSum*. The *digestFastqs()* metrics for the Li_tRNA_sel30 data set are averaged across the five runs on the single input sample, since only one run is required for *mutscan*. Total I/O volumes are separated in input and output, indicated above by I and O, respectively. RSS - resident set size.

While the evaluations mentioned above were all run with 10 cores, we also investigated how the key performance parameters scaled with the number of cores provided to *mutscan* when running the *digestFastqs()* function on a single pair of FASTQ files (Additional file 2:Figure S1). The results suggest that while the average load increases linearly with increasing number of cores, indicating good scalability of the parallel parts of the code, the benefit in terms of decreased total execution time is significantly reduced when more than 10 cores are used. This is likely explained by the constant runtime consumed by the serial parts in the code. In addition, for a fixed number of cores, the raw data processing performed by *digestFastqs()* scales linearly with the number of input reads in terms of execution time, and close to linearly in terms of memory requirement (Additional file 2:Figure S2).

### *mutscan* counts and fitness scores are comparable to those from *Enrich2* and *DiMSum*

In order to further compare the return values of *mutscan* to those obtained from *DiMSum* and *Enrich2*, we contrasted the sets of variants identified by the different tools. In addition, we chose a representative sample for each of the data sets, and contrasted the estimated counts for all variants identified by at least one tool (Figure 5, Additional file 2:Figure S3-S10). As previously described, we attempted to set the parameters of each tool in such a way as to maximize the similarity between the analysis workflows (see Methods). In general, most variants with up to two mutations (bases or amino acids, depending on the data set) were detected by all methods (Additional file 2:Figure S3, S5, S7, S9). The variants that were found with only one of the tools also generally had a lower abundance than the variants found consistently with all tools.

**Figure 5.**
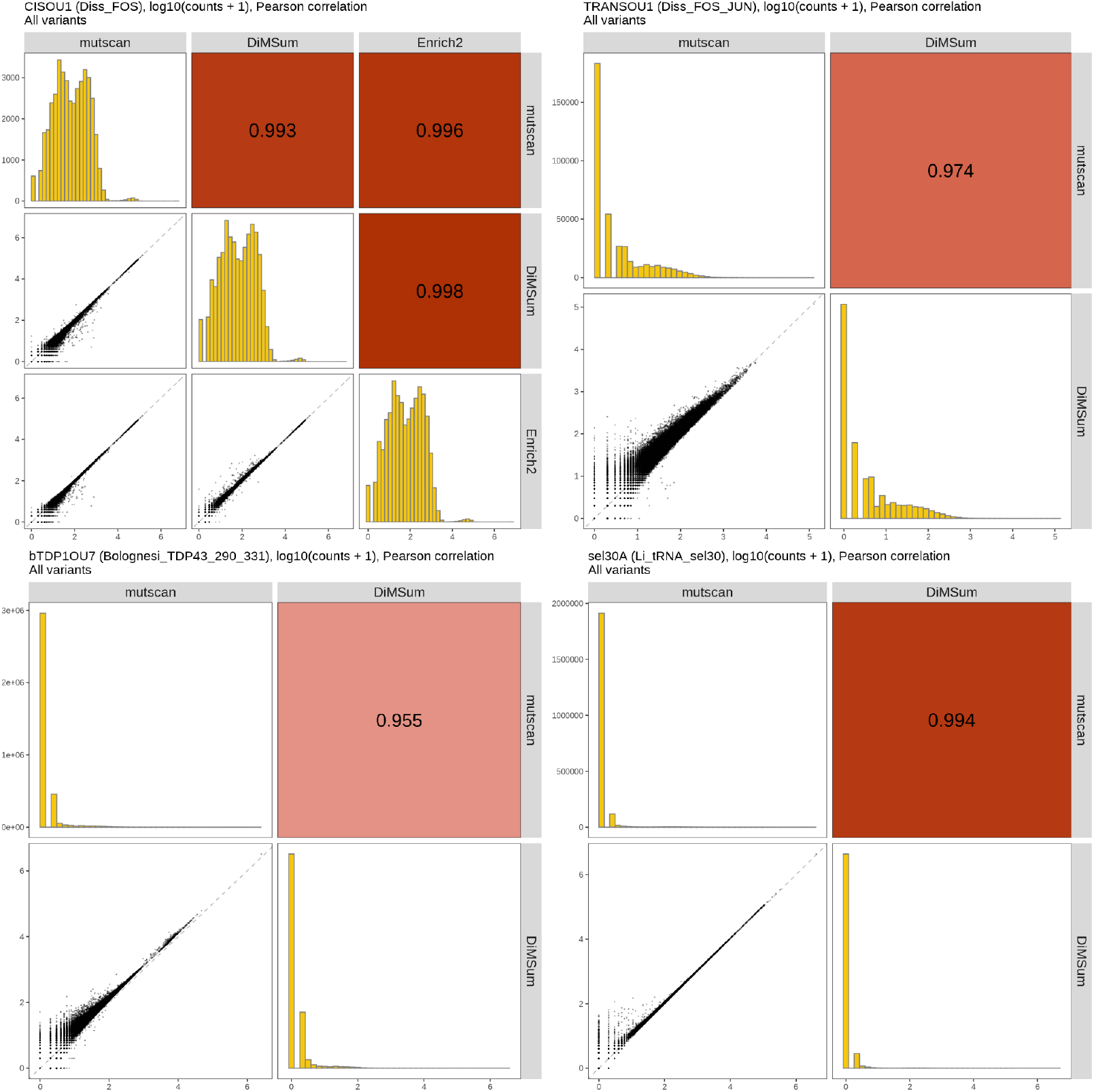
Comparison of counts estimated by *mutscan*, *DiMSum* and *Enrich2*. For each data set, a representative sample is shown (indicated in the respective figure titles, together with the data set). .

Also when comparing the estimated counts for the variants between the tools we noticed a high degree of similarity, but also data set-specific differences. For the Diss_FOS_JUN data set, we hypothesize that one reason for the generally higher counts observed with *DiMSum* than with *mutscan* is the different way in which the allowed codon mutations are specified. More precisely, with *mutscan* we are defining the format of any forbidden mutated codon (in our case, IUPAC code 'NNW'). *DiMSum*, on the other hand, approaches this by specifying the allowed IUPAC code for each position in the entire variable sequence. Hence, with this setup, mutated codons that already end in a 'T' or an 'A' in the wildtype sequence will not disqualify the read from being included in the analysis. The shift from the diagonal line in the Bolognesi_TDP43_290_331 data set may be caused by differences between approaches in *mutscan* and *DiMSum* for detecting the primer sequence immediately preceding the variable sequence. While *DiMSum* utilizes *cutadapt* to trim the unwanted sequence, *mutscan* requires a perfect match to the specified primer sequence whenever its position in the read is not fully specified.

Similarly to the sets of detected variants, the correlation between the estimated counts from the different tools also decreased as the number of mutations in the variant increased (Additional file 2:Figure S4, S6, S8, S10).

Finally, we calculated fitness scores using the three tools (Figure 6). For *mutscan*, we evaluated both built-in frameworks, based on *limma-voom* and *edgeR*, respectively. The log-fold change of a variant relative to that of the wildtype (obtained by using the latter as a replacement for the library size in the offset/normalization steps of *edgeR*/*limma*) is used as a fitness score for *mutscan*, and compared to the returned fitness score from *DiMSum*, and the average of the sample-wise scores from *Enrich2*. As for the variant counts, we see a strong correlation between the fitness scores from the different methods. This corroborates findings from (32), where fitness scores from *DiMSum* were found to be highly correlated with log-fold changes calculated using *DESeq2* (33) on the same count matrix. As for the detected variants, the correlation between the fitness scores is much stronger for variants with only a single mutation, and decreases as the number of mutations increases (Additional file 2:Figure S11-S14), possibly due to the lower absolute counts observed for variants with more mutations. It is worth noting that the dataset with the lowest correlation among the fitness scores (Bolognesi_TDP43_290_331) is also the one where the correlations between the *DiMSum* fitness scores for the individual replicates are the lowest (Additional file 2:Figure S15). Again, most of the discrepancies are contributed by variants with more mutated bases (Additional file 2:Figure S13).

**Figure 6.**
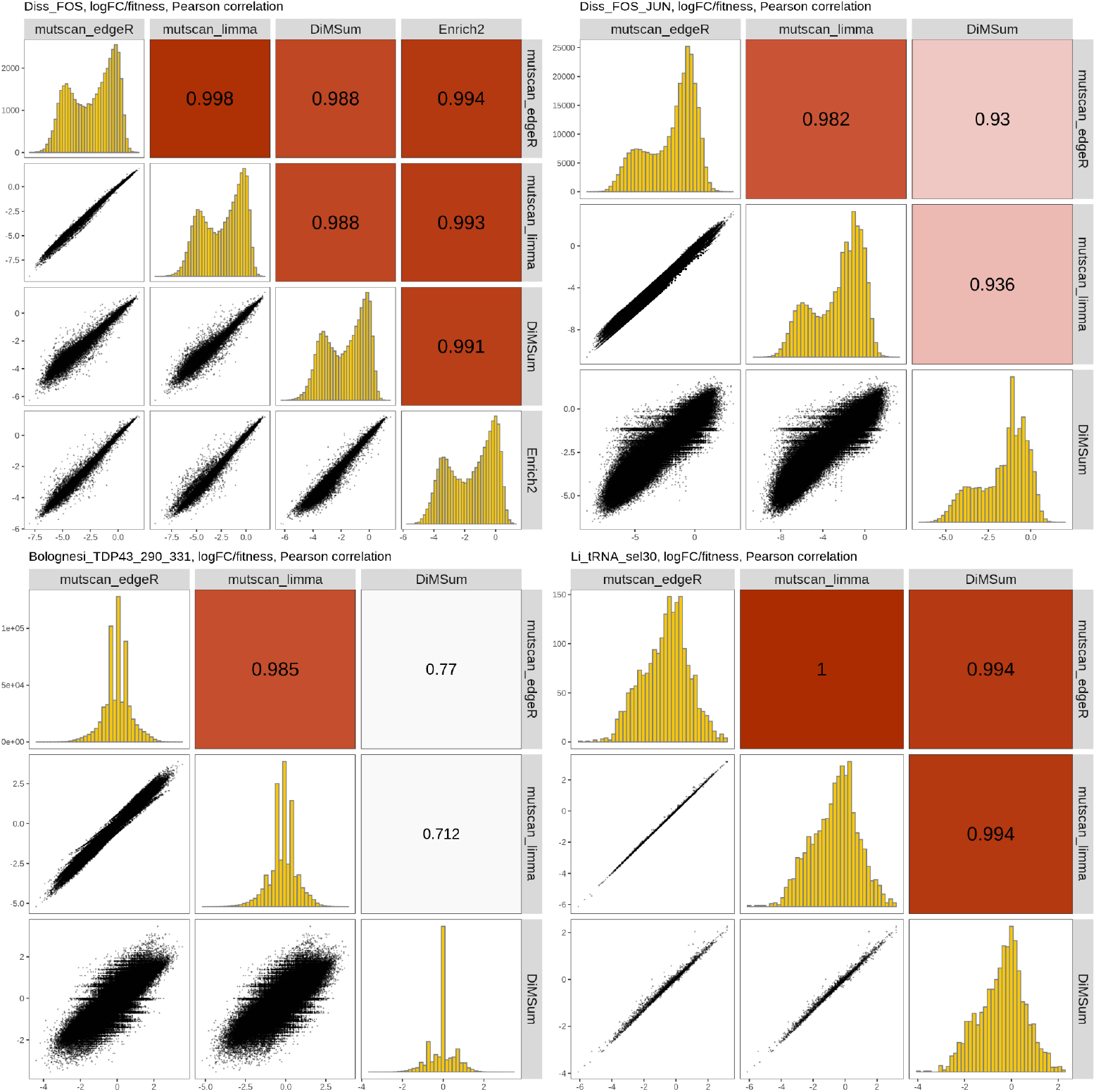
Comparison of fitness scores estimated by *mutscan*, *DiMSum* and *Enrich2*. For *mutscan*, the values are logFCs estimated by either *edgeR* or *limma*. For *DiMSum*, they correspond to the fitness score derived from all replicates. For *Enrich2*, they are the averages of the scores for the replicates.

## Conclusions

We have described *mutscan*, a flexible, easy-to-use R package for processing and statistical analysis of Multiplexed Assays of Variant Effect data. *mutscan* is designed in a modular way and is directly applicable also to other types of data aimed at identifying and tabulating substitution variants compared to a provided reference sequence, or tabulating unique sequences directly, potentially after collapsing variants within a certain distance. By leveraging established tools for statistical analysis of count data, the analytical framework provides a high degree of flexibility to address a variety of practical questions.

Since *mutscan* does not depend on external software libraries beyond R packages, it is easy to install across all major operating systems. At the same time it provides a high degree of interoperability, since the summarized data is represented in a *SummarizedExperiment* object, which can be directly used with a wide range of analysis and visualization functions within the Bioconductor ecosystem.

For many of the MAVE studies to date (including the ones used for evaluation in this study), the readout is the actual DNA sequence of the target protein(s) of interest. However, the field is increasingly moving towards instead sequencing unique barcodes associated with these variants, as this provides a way of better distinguishing true variants and sequencing errors, and also simplifies the analysis of longer protein sequences. *mutscan* supports also this type of experimental setup. If no reference sequence is provided, the variants (barcodes) will be represented by their sequence. Moreover, observed sequences can be collapsed if they are within a given distance, and have at least a pre-specified ratio of abundance in the sample. Current development work for *mutscan* includes streamlining the analysis workflow for large barcode sequencing experiments further, including the mapping of barcodes to true variants.

## Methods

Figure 1 provides an overview of the functionality implemented in *mutscan*. A typical workflow can often be summarized in three main steps: (1) processing of individual samples/sequencing libraries, (2) aggregation of the output from the individual samples into a single, combined object, (3) analysis and visualization.

### Processing of individual samples

The individual sample processing starts from a FASTQ file with sequencing reads (or a pair of FASTQ files for a paired-end sequencing experiment). Multiple (pairs of) FASTQ files per sample are supported. The *digestFastqs()* function takes these FASTQ files as input, and processes the reads to generate a variant count table. The first part of this analysis proceeds read by read, and consists of the following steps:

1. If applicable, search for user-specified adapter sequences and remove any read (pair) where these are detected.
2. Split the read (pair) into components as specified by the user (see Box 1 for details); in particular, extract the constant and variable parts of the reads. In the process, filter out reads that are not compatible with the user-specified composition.
3. Reverse complement the forward and/or reverse constant and variable sequences if requested.
4. If requested, merge the forward and reverse variable sequences. The user can specify minimum and maximum overlaps, minimum and maximum lengths of the merged sequence, and the maximal fraction of allowed mismatches. If no valid overlap is found satisfying these criteria, the read pair is filtered out.
5. Filter out reads where the average base quality in the variable sequence is below the user-specified threshold, or where the number of Ns in the variable or UMI sequence exceeds an imposed threshold.
6. If one or more wildtype/reference sequences are provided, compare the extracted variable region to these and find the closest match. If no wildtype sequence is found within an imposed mismatch limit, or if more than one wildtype sequence provides an equally good optimal match, the read (pair) is filtered out. Reads will also be filtered out if the base quality of the identified mutation(s) is below an imposed threshold, or if the mutated codon(s) matches a user-specified list of forbidden codons. Separate sets of wildtype sequences can be provided for the forward and reverse reads in a pair, if appropriate.
7. Similarly, compare the extracted constant sequences to provided reference sequences, and filter out reads where the difference exceeds a given threshold, or where more than one reference sequence provides equally good optimal matching.
8. If the read has not been filtered out in any of the steps above, store the observed sequence as well as an assigned 'mutant name', consisting of the name of the closest wildtype sequence together with the positions and sequences of the mutated nucleotides or codons. If no wildtype sequences are provided, the mutant name is the observed variable sequence.

The read processing is implemented in C++ for efficiency. Moreover, reads can be processed in parallel using OpenMP (34) in order to speed up this first step of an analysis. *mutscan* also supports writing all reads that are filtered out, together with the reason for the exclusion, to a FASTQ file (pair) for potential further processing/troubleshooting.

After all reads have been processed individually and the final set of sequences to retain has been determined, *mutscan* supports additional post-processing steps. If no wildtype sequences are provided, reads that are within a certain Hamming distance of each other can be collapsed (the assumption is that these correspond to sequencing or PCR error variants). This step will collapse a lower-frequency read to a higher-frequency one if their similarity as well as the abundance of the most frequent sequence and the ratio of the two abundances exceed given thresholds. The read counts for the collapsed sequences are summed, and all individual sequences contributing to a collapsed feature are recorded. Similarly, if the reads contain UMI sequences, these can be collapsed within a given variable sequence to avoid over-counting UMIs because of sequencing errors.

The individual sample processing with *digestFastqs()* returns an output object with four components. The count table records all the observed variants, together with their abundances (number of reads and/or UMIs), the number of mutated bases, codons and amino acids, and the set of observed sequences contributing to each variant. The filtering table summarizes the number of reads that were filtered out in each step outlined above. If a read is filtered out in one step, it will not be considered for the following ones. The output also contains a list of all argument values provided to *digestFastqs()*, as well as information about when the analysis was run, and with which version of *mutscan*. Finally, if one or more constant regions are included in the reads, *mutscan* tabulates the number of mismatches for each base quality, and returns the table.

### Merging processed data from multiple samples

A typical MAVE experiment consists of input and output (post-selection) libraries from multiple replicates. The process outlined in the previous section generates a count vector for each of these samples. To perform downstream analyses, the count vectors from the individual samples are merged into a single *SummarizedExperiment* object by the *summarizeExperiment()* function. Variants not detected in specific samples are thereby assigned a count of zero. In addition to the counts, the *summarizeExperiment* function propagates the filtering summary and parameter settings, as well as user-provided metadata about the samples, and stores these for convenient access in the joint object.

### Statistical analysis and visualization

In order to find variants that either increase or decrease their relative abundance upon selection, potentially compared to a wildtype variant, *mutscan* employs the widely used and established statistical models provided in the *edgeR* (29) and *limma* (35) Bioconductor packages to perform statistical analysis, leveraging the large number of distinct variants for improved inference. Normalization factors can be calculated using either the TMM method (36) (which generates sample-specific normalization factors) or the *csaw* package (37) (which generates feature- and sample-specific normalization factors). Alternatively, if the user wishes to calculate abundance changes relative to those of one or more wildtype sequences, the sum or geometric mean of the counts of the latter can be used as offsets. Similar approaches were previously explored, and found to perform well, for the analysis of multiple parallel reporter assays (38). The user can further select whether to use the *edgeR-QLF* (39) or the *limma-voom* (30) framework to fit a (generalized) linear model to each feature. The log-fold changes returned by these models can be used as fitness scores for downstream interpretation. In addition, for growth rate-based experiments, *mutscan* can be used to estimate PPI scores as described by (6).

In addition to the statistical analysis functionality, *mutscan* provides a variety of diagnostic plots, including a summary of the filtering steps, pairs plots displaying the correlations among samples, plots showing the distribution of abundances by sample, and static or interactive MA plots and volcano plots for easier interpretation of the statistical analysis results. It also provides a convenient wrapper function to generate a quality report in html format for an experiment.

### Comparison to *Enrich2* and *DiMSum*

To benchmark *mutscan*, we compared the computational performance metrics as well as the output counts and fitness scores to those from *DiMSum* (20) and *Enrich2* (19), which are both widely used tools for analysis of MAVE data. We aimed to set the parameters of the three methods in such a way that the output values were comparable, wherever possible. For the Diss_FOS data set, we limited the number of mutated nucleotides to 2, and analyzed the data on the nucleotide level rather than collapsing on the codon or amino acid level. We also did not use the information in the included UMIs, but counted the number of reads assigned to each variant. For the Bolognesi_TDP43_290_331 data set, we allowed up to three nucleotide substitutions. For the Li_tRNA_sel30 data set, we allowed any number of mutations, but filtered the quantified variants to only retain those with a count exceeding 2,000 in all the input samples, and exceeding 200 in all output samples. For the Diss_FOS_JUN data set, we instructed *mutscan* to allow up to one mutated codon in each of the two variable sequences, and only allowed mutated codons encoded by the 'NNS' IUPAC code. For *DiMSum*, we limited the total number of mutated amino acids to two (across the concatenated wild type sequence), indicated that the mutagenesis was done on the codon level, and provided the IUPAC code for the allowed nucleotide sequence. Before comparison, we further filtered out the *DiMSum* variants where the two mutated amino acids occurred in the same protein. We also removed variants with both non-synonymous and silent mutations from the *mutscan* output, as the default setting in *DiMSum* is to exclude these. Configuration files for all methods are available from https://github.com/fmicompbio/mutscan_manuscript_2022.

### Estimation of computational performance metrics

All analyses were run on a server with two Intel Xeon Platinum 8168 CPUs with a total of 48 cores, 1024 GB of random access memory and parallel file system accessed via GPFS. Memory requirement (max RSS), CPU usage, execution time and total I/O were measured using the benchmark directives of a slightly modified version of *snakemake* v7.8.3 (modified to return the values of read_chars and write_chars from *psutil*, in addition to the default read_bytes and write_bytes; patch file available at https://gist.github.com/mbstadler/3f5131b5aa88f87196d030a82081e1ea).

## Declarations

### Ethics approval and consent to participate

Not applicable

### Consent for publication

Not applicable

### Availability of data and materials

Project name: *mutscan* (v0.2.31)

Project home page: https://github.com/fmicompbio/mutscan, https://fmicompbio.github.io/mutscan/

Archived version: https://doi.org/10.5281/zenodo.7129132

Operating system(s): Platform independent

Programming language: R, C++

Other requirements: None

License: MIT

Any restrictions to use by non-academics: None

The code used to generate the results in the paper is available from https://github.com/fmicompbio/mutscan_manuscript_2022.

The data sets supporting the conclusions of this article are available in the Gene Expression Omnibus repository, accession numbers GSE102901 (https://www.ncbi.nlm.nih.gov/geo/query/acc.cgi?acc=GSE102901; Diss_FOS, diss_FOS_JUN), GSE128165 (https://www.ncbi.nlm.nih.gov/geo/query/acc.cgi?acc=GSE128165; Bolognesi_TDP43_290_331), GSE111508 (https://www.ncbi.nlm.nih.gov/geo/query/acc.cgi?acc=GSE111508; Li_tRNA_sel30).

### Authors’ contributions

CS: Conceptualization, Methodology, Software, Formal analysis, Writing-Original Draft, Visualization. AMB: Investigation, Writing-Original Draft, Validation. GD: Conceptualization, Resources, Validation, Writing - Review & Editing, Project administration. MBS: Conceptualization, Methodology, Software, Writing-Original Draft.

## Acknowledgements

The authors would like to thank current and former members of the Diss, Thomä and Stadler groups at FMI for discussions, testing and feedback.

