## Additional file 1 for "mutscan - a flexible R package for efficient end-to-end analysis of multiplexed assays of variant effect data": mutscan_AdditionalFile1.html

Additional file 1:Reprocessing FOS/JUN data from Diss & Lehner (2018)


Code 

- Show All Code
- Hide All Code
- Download Rmd

### Additional file 1:Reprocessing FOS/JUN data from Diss & Lehner (2018)

###### Charlotte Soneson, Alexandra M Bendel, Guillaume Diss, Michael B Stadler

### 1 Introduction

In this document, we reproduce the FOS/JUN protein-protein interaction analysis from the paper by Diss and Lehner (2018).
We assume that the FASTQ files have been downloaded from GEO record GSE102901 and are placed in a folder named `FASTQ`.
We start by tabulating the sample annotations, including the name we will use for downstream analysis (`Name`), the SRA ID (`SRAid`, for matching to the FASTQ files), the optical density (`OD`), the replicate number and the condition (whether it is an input or an output sample).
The goal will be to determine the fitness of each combination of FOS/JUN variants, relative to the wildtype sequence.

```
fastqdir <- "FASTQ"
list.files(fastqdir)
```

```
##  [1] "SRR5952429_1.fastq.gz" "SRR5952429_2.fastq.gz" "SRR5952430_1.fastq.gz"
##  [4] "SRR5952430_2.fastq.gz" "SRR5952431_1.fastq.gz" "SRR5952431_2.fastq.gz"
##  [7] "SRR5952432_1.fastq.gz" "SRR5952432_2.fastq.gz" "SRR5952433_1.fastq.gz"
## [10] "SRR5952433_2.fastq.gz" "SRR5952434_1.fastq.gz" "SRR5952434_2.fastq.gz"
```

```
(samples <- data.frame(
    Name = c("TRANSIN1", "TRANSIN2", "TRANSIN3", "TRANSOU1", "TRANSOU2", "TRANSOU3"),
    SRAid = c("SRR5952429", "SRR5952430", "SRR5952431", "SRR5952432", "SRR5952433", "SRR5952434"),
    OD = c(0.0025, 0.0025, 0.0025, 4.1762, 3.984, 3.9015),
    Replicate = c("R1", "R2", "R3", "R1", "R2", "R3"),
    Condition = c("IN", "IN", "IN", "OUT", "OUT", "OUT")
))
```

```
##       Name      SRAid     OD Replicate Condition
## 1 TRANSIN1 SRR5952429 0.0025        R1        IN
## 2 TRANSIN2 SRR5952430 0.0025        R2        IN
## 3 TRANSIN3 SRR5952431 0.0025        R3        IN
## 4 TRANSOU1 SRR5952432 4.1762        R1       OUT
## 5 TRANSOU2 SRR5952433 3.9840        R2       OUT
## 6 TRANSOU3 SRR5952434 3.9015        R3       OUT
```

### 2 Load required packages

```
suppressPackageStartupMessages({
    library(mutscan)
    library(SummarizedExperiment)
    library(scales)
    library(ggplot2)
    library(cowplot)
    library(GGally)
})
```

### 3 Process FASTQ files separately

The first step in the analysis workflow is to process the FASTQ files for the different samples separately, using the `digestFastqs()` function from the `mutscan` package.
In this case, the forward read contains a UMI, a constant sequence, and the sequence of the FOS variant.
Similarly, the reverse read contains a UMI, a constant sequence, and the sequence of the JUN variant.
As the two reads in a pair represent different protein variants, they don’t share any sequence and thus should not be merged for the purposes of the analysis.
By setting `mergeForwardReverse = FALSE`, we extract sequence components separately for the forward and reverse reads, and compare them to their respective wild type sequence.
The final mutant name will be a combination of the identified FOS and JUN mutations for the read pair.

```
res <- lapply(structure(samples$Name, names = samples$Name), function(s) {
    digestFastqs(fastqForward = file.path(fastqdir, paste0(samples$SRAid[samples$Name == s],
                                                           "_1.fastq.gz")),
                 fastqReverse = file.path(fastqdir, paste0(samples$SRAid[samples$Name == s],
                                                           "_2.fastq.gz")),
                 mergeForwardReverse = FALSE, 
                 adapterForward = "GGAAGAGCACACGTC",
                 adapterReverse = "GGAAGAGCGTCGTGT",
                 elementsForward = "SUCV",
                 elementLengthsForward = c(1, 10, 18, 96),
                 elementsReverse = "SUCV",
                 elementLengthsReverse = c(1, 8, 20, 96),
                 wildTypeForward = c(FOS = "ACTGATACACTCCAAGCGGAGACAGACCAACTAGAAGATGAGAAGTCTGCTTTGCAGACCGAGATTGCCAACCTGCTGAAGGAGAAGGAAAAACTA"),
                 wildTypeReverse = c(JUN = "ATCGCCCGGCTGGAGGAAAAAGTGAAAACCTTGAAAGCTCAGAACTCGGAGCTGGCGTCCACGGCCAACATGCTCAGGGAACAGGTGGCACAGCTT"),
                 constantForward = "AACCGGAGGAGGGAGCTG",
                 constantReverse = "GAAAAAGGAAGCTGGAGAGA",
                 nbrMutatedCodonsMaxForward = 1,
                 nbrMutatedCodonsMaxReverse = 1, 
                 forbiddenMutatedCodonsForward = "NNW",
                 forbiddenMutatedCodonsReverse = "NNW",
                 verbose = FALSE, 
                 nThreads = 10, 
                 maxNReads = -1)
})
```

The output of `digestFastqs()` is a list for each sample, containing the read and UMI counts for each variant, as well as a filtering summary.
For each variant, we also get the number of mutated bases, codons and amino acids, the type of mutations, and the nucleotide as well as amino acid sequence.
As we can see, the name assigned to each mutant combination consists of the name of the wildtype sequence(s), the codon position with a mutation (0 if no mutation is present), and the mutated codon sequence.
If we would like to focus on nucleotides rather than codons, that can be achieved by limiting the number of mutated bases rather than codons in the call to `digestFastqs()`.
In that case, the mutant name would consist of the wildtype sequence name, the position of the mutated nucleotide(s), and the observed nucleotide at that position.

```
## Our list has one entry per sample
names(res)
```

```
## [1] "TRANSIN1" "TRANSIN2" "TRANSIN3" "TRANSOU1" "TRANSOU2" "TRANSOU3"
```

```
## List entries for a specific sample
names(res$TRANSIN1)
```

```
## [1] "parameters"      "filterSummary"   "summaryTable"    "errorStatistics"
```

```
## Count table
head(res$TRANSIN1$summaryTable)
```

```
##           mutantName
## 1  FOS.0.WT_JUN.0.WT
## 2 FOS.0.WT_JUN.1.AAC
## 3 FOS.0.WT_JUN.1.AAG
## 4 FOS.0.WT_JUN.1.ACC
## 5 FOS.0.WT_JUN.1.ACG
## 6 FOS.0.WT_JUN.1.AGC
##                                                                                                                                                                                            sequence
## 1 ACTGATACACTCCAAGCGGAGACAGACCAACTAGAAGATGAGAAGTCTGCTTTGCAGACCGAGATTGCCAACCTGCTGAAGGAGAAGGAAAAACTA_ATCGCCCGGCTGGAGGAAAAAGTGAAAACCTTGAAAGCTCAGAACTCGGAGCTGGCGTCCACGGCCAACATGCTCAGGGAACAGGTGGCACAGCTT
## 2 ACTGATACACTCCAAGCGGAGACAGACCAACTAGAAGATGAGAAGTCTGCTTTGCAGACCGAGATTGCCAACCTGCTGAAGGAGAAGGAAAAACTA_AACGCCCGGCTGGAGGAAAAAGTGAAAACCTTGAAAGCTCAGAACTCGGAGCTGGCGTCCACGGCCAACATGCTCAGGGAACAGGTGGCACAGCTT
## 3 ACTGATACACTCCAAGCGGAGACAGACCAACTAGAAGATGAGAAGTCTGCTTTGCAGACCGAGATTGCCAACCTGCTGAAGGAGAAGGAAAAACTA_AAGGCCCGGCTGGAGGAAAAAGTGAAAACCTTGAAAGCTCAGAACTCGGAGCTGGCGTCCACGGCCAACATGCTCAGGGAACAGGTGGCACAGCTT
## 4 ACTGATACACTCCAAGCGGAGACAGACCAACTAGAAGATGAGAAGTCTGCTTTGCAGACCGAGATTGCCAACCTGCTGAAGGAGAAGGAAAAACTA_ACCGCCCGGCTGGAGGAAAAAGTGAAAACCTTGAAAGCTCAGAACTCGGAGCTGGCGTCCACGGCCAACATGCTCAGGGAACAGGTGGCACAGCTT
## 5 ACTGATACACTCCAAGCGGAGACAGACCAACTAGAAGATGAGAAGTCTGCTTTGCAGACCGAGATTGCCAACCTGCTGAAGGAGAAGGAAAAACTA_ACGGCCCGGCTGGAGGAAAAAGTGAAAACCTTGAAAGCTCAGAACTCGGAGCTGGCGTCCACGGCCAACATGCTCAGGGAACAGGTGGCACAGCTT
## 6 ACTGATACACTCCAAGCGGAGACAGACCAACTAGAAGATGAGAAGTCTGCTTTGCAGACCGAGATTGCCAACCTGCTGAAGGAGAAGGAAAAACTA_AGCGCCCGGCTGGAGGAAAAAGTGAAAACCTTGAAAGCTCAGAACTCGGAGCTGGCGTCCACGGCCAACATGCTCAGGGAACAGGTGGCACAGCTT
##   nbrReads maxNbrReads nbrUmis nbrMutBases nbrMutCodons nbrMutAAs varLengths
## 1    16680       16680   16649           0            0         0      96_96
## 2      285         285     284           1            1         1      96_96
## 3      382         382     381           2            1         1      96_96
## 4      408         408     407           1            1         1      96_96
## 5      254         254     254           2            1         1      96_96
## 6      179         179     179           1            1         1      96_96
##        mutantNameAA mutationTypes
## 1 FOS.0.WT_JUN.0.WT              
## 2  FOS.0.WT_JUN.1.N nonsynonymous
## 3  FOS.0.WT_JUN.1.K nonsynonymous
## 4  FOS.0.WT_JUN.1.T nonsynonymous
## 5  FOS.0.WT_JUN.1.T nonsynonymous
## 6  FOS.0.WT_JUN.1.S nonsynonymous
##                                                          sequenceAA
## 1 TDTLQAETDQLEDEKSALQTEIANLLKEKEKL_IARLEEKVKTLKAQNSELASTANMLREQVAQL
## 2 TDTLQAETDQLEDEKSALQTEIANLLKEKEKL_NARLEEKVKTLKAQNSELASTANMLREQVAQL
## 3 TDTLQAETDQLEDEKSALQTEIANLLKEKEKL_KARLEEKVKTLKAQNSELASTANMLREQVAQL
## 4 TDTLQAETDQLEDEKSALQTEIANLLKEKEKL_TARLEEKVKTLKAQNSELASTANMLREQVAQL
## 5 TDTLQAETDQLEDEKSALQTEIANLLKEKEKL_TARLEEKVKTLKAQNSELASTANMLREQVAQL
## 6 TDTLQAETDQLEDEKSALQTEIANLLKEKEKL_SARLEEKVKTLKAQNSELASTANMLREQVAQL
```

```
## Filter summary
res$TRANSIN1$filterSummary
```

```
##   nbrTotal f1_nbrAdapter f2_nbrNoPrimer f3_nbrReadWrongLength
## 1 37117489      12157899              0                     0
##   f4_nbrNoValidOverlap f5_nbrAvgVarQualTooLow f6_nbrTooManyNinVar
## 1                    0                 152312             1783532
##   f7_nbrTooManyNinUMI f8_nbrTooManyBestWTHits f9_nbrMutQualTooLow
## 1                1111                       0                   0
##   f10a_nbrTooManyMutCodons f10b_nbrTooManyMutBases f11_nbrForbiddenCodons
## 1                 12033891                       0                 193216
##   f12_nbrTooManyMutConstant f13_nbrTooManyBestConstantHits nbrRetained
## 1                         0                              0    10795528
```

```
## Parameters used for the processing, including mutscan version and analysis date
res$TRANSIN1$parameters$processingInfo
```

```
## [1] "Processed by mutscan v0.2.31 on 2022-09-30 20:04:21"
```

Since we provided a constant sequence for our experiment, `mutscan` additionally outputs a table that lets us estimate the sequencing error rate.

```
(propErrorsConstantF <- sum(res$TRANSIN1$errorStatistics$nbrMismatchForward) /
   (nchar(res$TRANSIN1$parameters$constantForward) * res$TRANSIN1$filterSummary$nbrRetained))
```

```
## [1] 0.001811887
```

```
(propErrorsConstantR <- sum(res$TRANSIN1$errorStatistics$nbrMismatchReverse) /
   (nchar(res$TRANSIN1$parameters$constantReverse) * res$TRANSIN1$filterSummary$nbrRetained))
```

```
## [1] 0.006168948
```

### 4 Summarizing output from all samples

While the `digestFastqs()` output for each sample can be explored directly, for convenience we merge the data from all samples into a single SummarizedExperiment object.
This object contains all information from the `digestFastqs()` output above, as well as the information from the sample metadata table.

```
samples
```

```
##       Name      SRAid     OD Replicate Condition
## 1 TRANSIN1 SRR5952429 0.0025        R1        IN
## 2 TRANSIN2 SRR5952430 0.0025        R2        IN
## 3 TRANSIN3 SRR5952431 0.0025        R3        IN
## 4 TRANSOU1 SRR5952432 4.1762        R1       OUT
## 5 TRANSOU2 SRR5952433 3.9840        R2       OUT
## 6 TRANSOU3 SRR5952434 3.9015        R3       OUT
```

```
se <- summarizeExperiment(res, 
                          coldata = samples,
                          countType = "umis")
se
```

```
## class: SummarizedExperiment 
## dim: 964285 6 
## metadata(3): parameters countType mutNameDelimiter
## assays(1): counts
## rownames(964285): FOS.0.WT_JUN.0.WT FOS.0.WT_JUN.1.AAC ...
##   FOS.9.TTG_JUN.9.TTC FOS.9.TTG_JUN.9.TTG
## rowData names(15): mutantName sequence ... mutationTypes varLengths
## colnames(6): TRANSIN1 TRANSIN2 ... TRANSOU2 TRANSOU3
## colData names(21): Name SRAid ... f13_nbrTooManyBestConstantHits
##   nbrRetained
```

```
## Variant information
rowData(se)
```

```
## DataFrame with 964285 rows and 15 columns
##                              mutantName               sequence nbrMutBases
##                             <character>            <character> <character>
## FOS.0.WT_JUN.0.WT     FOS.0.WT_JUN.0.WT ACTGATACACTCCAAGCGGA..           0
## FOS.0.WT_JUN.1.AAC   FOS.0.WT_JUN.1.AAC ACTGATACACTCCAAGCGGA..           1
## FOS.0.WT_JUN.1.AAG   FOS.0.WT_JUN.1.AAG ACTGATACACTCCAAGCGGA..           2
## FOS.0.WT_JUN.1.ACC   FOS.0.WT_JUN.1.ACC ACTGATACACTCCAAGCGGA..           1
## FOS.0.WT_JUN.1.ACG   FOS.0.WT_JUN.1.ACG ACTGATACACTCCAAGCGGA..           2
## ...                                 ...                    ...         ...
## FOS.9.TTG_JUN.9.TCG FOS.9.TTG_JUN.9.TCG ACTGATACACTCCAAGCGGA..           6
## FOS.9.TTG_JUN.9.TGC FOS.9.TTG_JUN.9.TGC ACTGATACACTCCAAGCGGA..           6
## FOS.9.TTG_JUN.9.TGG FOS.9.TTG_JUN.9.TGG ACTGATACACTCCAAGCGGA..           6
## FOS.9.TTG_JUN.9.TTC FOS.9.TTG_JUN.9.TTC ACTGATACACTCCAAGCGGA..           6
## FOS.9.TTG_JUN.9.TTG FOS.9.TTG_JUN.9.TTG ACTGATACACTCCAAGCGGA..           6
##                     minNbrMutBases maxNbrMutBases nbrMutCodons minNbrMutCodons
##                          <integer>      <integer>  <character>       <integer>
## FOS.0.WT_JUN.0.WT                0              0            0               0
## FOS.0.WT_JUN.1.AAC               1              1            1               1
## FOS.0.WT_JUN.1.AAG               2              2            1               1
## FOS.0.WT_JUN.1.ACC               1              1            1               1
## FOS.0.WT_JUN.1.ACG               2              2            1               1
## ...                            ...            ...          ...             ...
## FOS.9.TTG_JUN.9.TCG              6              6            2               2
## FOS.9.TTG_JUN.9.TGC              6              6            2               2
## FOS.9.TTG_JUN.9.TGG              6              6            2               2
## FOS.9.TTG_JUN.9.TTC              6              6            2               2
## FOS.9.TTG_JUN.9.TTG              6              6            2               2
##                     maxNbrMutCodons   nbrMutAAs minNbrMutAAs maxNbrMutAAs
##                           <integer> <character>    <integer>    <integer>
## FOS.0.WT_JUN.0.WT                 0           0            0            0
## FOS.0.WT_JUN.1.AAC                1           1            1            1
## FOS.0.WT_JUN.1.AAG                1           1            1            1
## FOS.0.WT_JUN.1.ACC                1           1            1            1
## FOS.0.WT_JUN.1.ACG                1           1            1            1
## ...                             ...         ...          ...          ...
## FOS.9.TTG_JUN.9.TCG               2           2            2            2
## FOS.9.TTG_JUN.9.TGC               2           2            2            2
## FOS.9.TTG_JUN.9.TGG               2           2            2            2
## FOS.9.TTG_JUN.9.TTC               2           2            2            2
## FOS.9.TTG_JUN.9.TTG               2           2            2            2
##                                 sequenceAA      mutantNameAA mutationTypes
##                                <character>       <character>   <character>
## FOS.0.WT_JUN.0.WT   TDTLQAETDQLEDEKSALQT.. FOS.0.WT_JUN.0.WT              
## FOS.0.WT_JUN.1.AAC  TDTLQAETDQLEDEKSALQT..  FOS.0.WT_JUN.1.N nonsynonymous
## FOS.0.WT_JUN.1.AAG  TDTLQAETDQLEDEKSALQT..  FOS.0.WT_JUN.1.K nonsynonymous
## FOS.0.WT_JUN.1.ACC  TDTLQAETDQLEDEKSALQT..  FOS.0.WT_JUN.1.T nonsynonymous
## FOS.0.WT_JUN.1.ACG  TDTLQAETDQLEDEKSALQT..  FOS.0.WT_JUN.1.T nonsynonymous
## ...                                    ...               ...           ...
## FOS.9.TTG_JUN.9.TCG TDTLQAETLQLEDEKSALQT..   FOS.9.L_JUN.9.S nonsynonymous
## FOS.9.TTG_JUN.9.TGC TDTLQAETLQLEDEKSALQT..   FOS.9.L_JUN.9.C nonsynonymous
## FOS.9.TTG_JUN.9.TGG TDTLQAETLQLEDEKSALQT..   FOS.9.L_JUN.9.W nonsynonymous
## FOS.9.TTG_JUN.9.TTC TDTLQAETLQLEDEKSALQT..   FOS.9.L_JUN.9.F nonsynonymous
## FOS.9.TTG_JUN.9.TTG TDTLQAETLQLEDEKSALQT..   FOS.9.L_JUN.9.L nonsynonymous
##                      varLengths
##                     <character>
## FOS.0.WT_JUN.0.WT         96_96
## FOS.0.WT_JUN.1.AAC        96_96
## FOS.0.WT_JUN.1.AAG        96_96
## FOS.0.WT_JUN.1.ACC        96_96
## FOS.0.WT_JUN.1.ACG        96_96
## ...                         ...
## FOS.9.TTG_JUN.9.TCG       96_96
## FOS.9.TTG_JUN.9.TGC       96_96
## FOS.9.TTG_JUN.9.TGG       96_96
## FOS.9.TTG_JUN.9.TTC       96_96
## FOS.9.TTG_JUN.9.TTG       96_96
```

```
## Sample information
colData(se)
```

```
## DataFrame with 6 rows and 21 columns
##                 Name       SRAid        OD   Replicate   Condition  nbrTotal
##          <character> <character> <numeric> <character> <character> <integer>
## TRANSIN1    TRANSIN1  SRR5952429    0.0025          R1          IN  37117489
## TRANSIN2    TRANSIN2  SRR5952430    0.0025          R2          IN  34097299
## TRANSIN3    TRANSIN3  SRR5952431    0.0025          R3          IN  37640983
## TRANSOU1    TRANSOU1  SRR5952432    4.1762          R1         OUT  41662789
## TRANSOU2    TRANSOU2  SRR5952433    3.9840          R2         OUT  42541479
## TRANSOU3    TRANSOU3  SRR5952434    3.9015          R3         OUT  52616569
##          f1_nbrAdapter f2_nbrNoPrimer f3_nbrReadWrongLength
##              <integer>      <integer>             <integer>
## TRANSIN1      12157899              0                     0
## TRANSIN2       4315856              0                     0
## TRANSIN3      10230180              0                     0
## TRANSOU1      16139824              0                     0
## TRANSOU2      12757387              0                     0
## TRANSOU3      26030172              0                     0
##          f4_nbrNoValidOverlap f5_nbrAvgVarQualTooLow f6_nbrTooManyNinVar
##                     <integer>              <integer>           <integer>
## TRANSIN1                    0                 152312             1783532
## TRANSIN2                    0                 183561             2112765
## TRANSIN3                    0                 185871             1958471
## TRANSOU1                    0                 204382             1842382
## TRANSOU2                    0                 258212             2123540
## TRANSOU3                    0                 270528             1946686
##          f7_nbrTooManyNinUMI f8_nbrTooManyBestWTHits f9_nbrMutQualTooLow
##                    <integer>               <integer>           <integer>
## TRANSIN1                1111                       0                   0
## TRANSIN2                1391                       0                   0
## TRANSIN3                1190                       0                   0
## TRANSOU1                1047                       0                   0
## TRANSOU2                1327                       0                   0
## TRANSOU3                1148                       0                   0
##          f10a_nbrTooManyMutCodons f10b_nbrTooManyMutBases
##                         <integer>               <integer>
## TRANSIN1                 12033891                       0
## TRANSIN2                 13287986                       0
## TRANSIN3                 13134347                       0
## TRANSOU1                 11412147                       0
## TRANSOU2                 12952351                       0
## TRANSOU3                 11863729                       0
##          f11_nbrForbiddenCodons f12_nbrTooManyMutConstant
##                       <integer>                 <integer>
## TRANSIN1                 193216                         0
## TRANSIN2                 221384                         0
## TRANSIN3                 203401                         0
## TRANSOU1                 235187                         0
## TRANSOU2                 271692                         0
## TRANSOU3                 238182                         0
##          f13_nbrTooManyBestConstantHits nbrRetained
##                               <integer>   <integer>
## TRANSIN1                              0    10795528
## TRANSIN2                              0    13974356
## TRANSIN3                              0    11927523
## TRANSOU1                              0    11827820
## TRANSOU2                              0    14176970
## TRANSOU3                              0    12266124
```

### 5 Diagnostic plots

At this point, we can look at some diagnostic plots.
First, we display the number of identified variants with a given number of base mutations.

```
(g1 <- ggplot(as.data.frame(rowData(se)), aes(x = minNbrMutBases)) + 
     geom_bar() + theme_bw() + 
     scale_y_log10() + 
     labs(x = "Number of mutated bases (FOS + JUN)",
          y = "Number of variants") + 
     theme(axis.text = element_text(size = 15),
           axis.title = element_text(size = 15)))
```

We can also visualize the average number of counts (across all samples) for variants with different numbers of mutations.

```
df <- data.frame(nbrmut = factor(rowData(se)$minNbrMutBases),
                 avelogcount = rowMeans(log10(assay(se, "counts") + 1)))
(g2 <- ggplot(df, aes(x = nbrmut, y = avelogcount)) + 
        geom_violin() + 
        stat_summary(fun = "median",
                     geom = "crossbar", 
                     width = 0.25,
                     color = "red") + 
        geom_point(data = df %>% dplyr::filter(nbrmut == 0), size = 3, color = "black") + 
        theme_bw() + 
        labs(x = "Number of mutated bases (FOS + JUN)", y = "Mean (log10(count + 1))") + 
        theme(axis.text = element_text(size = 15),
              axis.title = element_text(size = 15)))
```

```
## Warning: Groups with fewer than two data points have been dropped.
```

From the plots above, we conclude that there are many more different variants with a large number of base mutations, but each of them is considerably less abundant than variants with fewer mutations.

Next, we display the number of reads that remain after each of the filtering criteria in `digestFastqs()`, for each of the samples, as well as the fraction of reads filtered out by each of the criteria.

```
(g3 <- plotFiltering(se, valueType = "reads", onlyActiveFilters = TRUE, 
                     plotType = "remaining", facetBy = "sample", numberSize = 3))
```

```
(g4 <- plotFiltering(se, valueType = "fractions", onlyActiveFilters = TRUE,
                     plotType = "filtered", facetBy = "step", numberSize = 3))
```

### 6 Concordance among the samples

In addition to sample-specific diagnostic plots like the ones shown above, we can also look at the concordance among the samples, in terms of the correlation between their count vectors.

```
(g5 <- plotPairs(se, selAssay = "counts", addIdentityLine = TRUE))
```

### 7 Collapsing data by amino acid

In the objects that we have been working with so far, each row corresponds to a distinct variant (codon mutation).
In some cases, we would like to collapse variants with the same amino acid sequence.
This can be done using the `collapseMutantsByAA()` function, which returns another SummarizedExperiment, where each row represents a specific amino acid sequence.

```
se <- collapseMutantsByAA(se)
```

We can see that this object is smaller than the previous one (fewer rows), as expected.
In addition, each row now corresponds to potentially several different nucleotide sequences and number of mutated bases.
For example, the “wild type” amino acid sequence is represented by individual sequences with up to six silent mutations.

```
se
```

```
## class: SummarizedExperiment 
## dim: 401200 6 
## metadata(3): parameters countType mutNameDelimiter
## assays(1): counts
## rownames(401200): FOS.0.WT_JUN.0.WT FOS.0.WT_JUN.1.* ...
##   FOS.9.Y_JUN.9.W FOS.9.Y_JUN.9.Y
## rowData names(14): mutantNameAA sequence ... maxNbrMutCodons
##   maxNbrMutAAs
## colnames(6): TRANSIN1 TRANSIN2 ... TRANSOU2 TRANSOU3
## colData names(21): Name SRAid ... f13_nbrTooManyBestConstantHits
##   nbrRetained
```

```
rowData(se)
```

```
## DataFrame with 401200 rows and 14 columns
##                        mutantNameAA               sequence
##                         <character>            <character>
## FOS.0.WT_JUN.0.WT FOS.0.WT_JUN.0.WT ACCGATACACTCCAAGCGGA..
## FOS.0.WT_JUN.1.*   FOS.0.WT_JUN.1.* ACTGACACACTCCAAGCGGA..
## FOS.0.WT_JUN.1.A   FOS.0.WT_JUN.1.A ACCGATACACTCCAAGCGGA..
## FOS.0.WT_JUN.1.C   FOS.0.WT_JUN.1.C ACCGATACACTCCAAGCGGA..
## FOS.0.WT_JUN.1.D   FOS.0.WT_JUN.1.D ACCGATACACTCCAAGCGGA..
## ...                             ...                    ...
## FOS.9.Y_JUN.9.S     FOS.9.Y_JUN.9.S ACTGATACACTCCAAGCGGA..
## FOS.9.Y_JUN.9.T     FOS.9.Y_JUN.9.T ACTGATACACTCCAAGCGGA..
## FOS.9.Y_JUN.9.V     FOS.9.Y_JUN.9.V ACTGATACACTCCAAGCGGA..
## FOS.9.Y_JUN.9.W     FOS.9.Y_JUN.9.W ACTGATACACTCCAAGCGGA..
## FOS.9.Y_JUN.9.Y     FOS.9.Y_JUN.9.Y ACTGATACACTCCAAGCGGA..
##                               mutantName             sequenceAA
##                              <character>            <character>
## FOS.0.WT_JUN.0.WT FOS.0.WT_JUN.0.WT,FO.. TDTLQAETDQLEDEKSALQT..
## FOS.0.WT_JUN.1.*  FOS.0.WT_JUN.1.TAG,F.. TDTLQAETDQLEDEKSALQT..
## FOS.0.WT_JUN.1.A  FOS.0.WT_JUN.1.GCC,F.. TDTLQAETDQLEDEKSALQT..
## FOS.0.WT_JUN.1.C  FOS.0.WT_JUN.1.TGC,F.. TDTLQAETDQLEDEKSALQT..
## FOS.0.WT_JUN.1.D  FOS.0.WT_JUN.1.GAC,F.. TDTLQAETDQLEDEKSALQT..
## ...                                  ...                    ...
## FOS.9.Y_JUN.9.S   FOS.9.TAC_JUN.9.AGC,.. TDTLQAETYQLEDEKSALQT..
## FOS.9.Y_JUN.9.T   FOS.9.TAC_JUN.9.ACC,.. TDTLQAETYQLEDEKSALQT..
## FOS.9.Y_JUN.9.V   FOS.9.TAC_JUN.9.GTC,.. TDTLQAETYQLEDEKSALQT..
## FOS.9.Y_JUN.9.W      FOS.9.TAC_JUN.9.TGG TDTLQAETYQLEDEKSALQT..
## FOS.9.Y_JUN.9.Y      FOS.9.TAC_JUN.9.TAC TDTLQAETYQLEDEKSALQT..
##                          mutationTypes   nbrMutBases nbrMutCodons   nbrMutAAs
##                            <character>   <character>  <character> <character>
## FOS.0.WT_JUN.0.WT               silent 0,1,2,3,4,5,6        0,1,2           0
## FOS.0.WT_JUN.1.*           silent,stop       3,4,5,6          1,2           1
## FOS.0.WT_JUN.1.A  nonsynonymous,silent     2,3,4,5,6          1,2           1
## FOS.0.WT_JUN.1.C  nonsynonymous,silent       2,3,4,5          1,2           1
## FOS.0.WT_JUN.1.D  nonsynonymous,silent       2,3,4,5          1,2           1
## ...                                ...           ...          ...         ...
## FOS.9.Y_JUN.9.S          nonsynonymous           3,4            2           2
## FOS.9.Y_JUN.9.T          nonsynonymous             3            2           2
## FOS.9.Y_JUN.9.V          nonsynonymous             4            2           2
## FOS.9.Y_JUN.9.W          nonsynonymous             4            2           2
## FOS.9.Y_JUN.9.Y          nonsynonymous             3            2           2
##                   minNbrMutBases minNbrMutCodons minNbrMutAAs maxNbrMutBases
##                        <integer>       <integer>    <integer>      <integer>
## FOS.0.WT_JUN.0.WT              0               0            0              6
## FOS.0.WT_JUN.1.*               3               1            1              6
## FOS.0.WT_JUN.1.A               2               1            1              6
## FOS.0.WT_JUN.1.C               2               1            1              5
## FOS.0.WT_JUN.1.D               2               1            1              5
## ...                          ...             ...          ...            ...
## FOS.9.Y_JUN.9.S                3               2            2              4
## FOS.9.Y_JUN.9.T                3               2            2              3
## FOS.9.Y_JUN.9.V                4               2            2              4
## FOS.9.Y_JUN.9.W                4               2            2              4
## FOS.9.Y_JUN.9.Y                3               2            2              3
##                   maxNbrMutCodons maxNbrMutAAs
##                         <integer>    <integer>
## FOS.0.WT_JUN.0.WT               2            0
## FOS.0.WT_JUN.1.*                2            1
## FOS.0.WT_JUN.1.A                2            1
## FOS.0.WT_JUN.1.C                2            1
## FOS.0.WT_JUN.1.D                2            1
## ...                           ...          ...
## FOS.9.Y_JUN.9.S                 2            2
## FOS.9.Y_JUN.9.T                 2            2
## FOS.9.Y_JUN.9.V                 2            2
## FOS.9.Y_JUN.9.W                 2            2
## FOS.9.Y_JUN.9.Y                 2            2
```

### 8 Calculating fitness scores

`mutscan` provides multiple options for calculating fitness scores.
For growth rate-based assays, the fitness score as defined by Diss and Lehner (2018) can be calculated for each sample using the `calculateFitnessScore()` function.

```
ppis <- calculateFitnessScore(se = se, pairingCol = "Replicate", 
                              ODCols = c("OD"),
                              comparison = c("Condition", "OUT", "IN"),
                              WTrows = "FOS.0.WT_JUN.0.WT")
head(ppis[order(abs(rowMeans(ppis)), decreasing = TRUE), , drop = FALSE])
```

```
##                   OUT_vs_IN_replR1 OUT_vs_IN_replR2 OUT_vs_IN_replR3
## FOS.8.I_JUN.5.L           1.257708         1.177871         1.184341
## FOS.16.E_JUN.17.V         1.275183         1.156391         1.184341
## FOS.8.I_JUN.5.R           1.271891         1.149459         1.150909
## FOS.8.F_JUN.3.T           1.121530         1.198412         1.240593
## FOS.1.L_JUN.10.H          1.113180         1.275265         1.163643
## FOS.7.L_JUN.5.C           1.253877         1.177871         1.111315
```

```
## The wildtype sequence has a fitness score of 1, by construction
ppis["FOS.0.WT_JUN.0.WT", , drop = FALSE]
```

```
##                   OUT_vs_IN_replR1 OUT_vs_IN_replR2 OUT_vs_IN_replR3
## FOS.0.WT_JUN.0.WT                1                1                1
```

In addition, and more generally, a fitness score can be calculated as the log-fold change of output vs input samples (paired by the Replicate column), either in absolute terms or relative to the wildtype sequence (by specifying the `WTrows` argument to the `calculateRelativeFC()` function).
`mutscan` allows using either edgeR or limma-voom for calculation of log-fold changes.

```
edger_scores <- calculateRelativeFC(
    se = se,
    design = model.matrix(~ Replicate + Condition,
                          data = colData(se)),
    coef = "ConditionOUT", pseudocount = 1,
    WTrows = "FOS.0.WT_JUN.0.WT",
    method = "edgeR")
head(edger_scores[order(edger_scores$PValue), , drop = FALSE])
```

```
##                       logFC   logCPM         F PValue FDR logFC_shrunk df.total
## FOS.0.WT_JUN.1.G  -1.011928 8.844134  2839.950      0   0    -1.011790   802400
## FOS.0.WT_JUN.1.P  -1.345886 7.430130  1907.883      0   0    -1.345400   802400
## FOS.0.WT_JUN.10.* -6.410407 4.014574  1831.750      0   0    -6.284890   802400
## FOS.0.WT_JUN.10.P -5.276403 6.861820 13134.671      0   0    -5.267873   802400
## FOS.0.WT_JUN.11.A -3.239381 5.969672  3504.138      0   0    -3.235016   802400
## FOS.0.WT_JUN.11.C -3.961166 4.692851  1870.035      0   0    -3.944772   802400
##                   df.prior df.test
## FOS.0.WT_JUN.1.G       Inf       1
## FOS.0.WT_JUN.1.P       Inf       1
## FOS.0.WT_JUN.10.*      Inf       1
## FOS.0.WT_JUN.10.P      Inf       1
## FOS.0.WT_JUN.11.A      Inf       1
## FOS.0.WT_JUN.11.C      Inf       1
```

```
## The wildtype sequence has a log-fold change close to 0, by construction
edger_scores["FOS.0.WT_JUN.0.WT", , drop = FALSE]
```

```
##                           logFC   logCPM            F    PValue FDR
## FOS.0.WT_JUN.0.WT -2.068241e-15 13.06047 3.295153e-09 0.9999975   1
##                   logFC_shrunk df.total df.prior df.test
## FOS.0.WT_JUN.0.WT 4.905382e-17   802400      Inf       1
```

### 9 Reproducing some figures from Diss and Lehner (2018)

In this section, we use the results generated above to reproduce some of the figures from Diss and Lehner (2018).
First, we filter the variants and keep only those with at least 11 counts in all input samples, and at least 1 count in all output samples.
We also remove all variants with a premature stop codon.

```
keepVariants <- 
    rownames(se)[
        DelayedArray::rowMins(as.matrix(assay(se[, se$Condition == "IN"],
                                              "counts"))) > 10 & 
            DelayedArray::rowMins(as.matrix(assay(se[, se$Condition == "OUT"],
                                                  "counts"))) > 0]
keepVariants <- setdiff(keepVariants, grep("*", keepVariants, fixed = TRUE, value = TRUE))
length(keepVariants)
```

```
## [1] 108841
```

We also summarize the PPI scores and log-fold changes in a table, for convenience.

```
df0 <- data.frame(mutation = rownames(ppis), 
                  PPI1 = ppis[, "OUT_vs_IN_replR1"],
                  PPI2 = ppis[, "OUT_vs_IN_replR2"],
                  PPI3 = ppis[, "OUT_vs_IN_replR3"],
                  avePPI = rowMeans(ppis, na.rm = TRUE),
                  stringsAsFactors = FALSE) %>%
    dplyr::full_join(as.data.frame(edger_scores) %>% 
                         tibble::rownames_to_column("mutation") %>%
                         dplyr::select(mutation, logFC_shrunk, logCPM, FDR),
                     by = "mutation") %>%
    dplyr::mutate(keepVariants = mutation %in% keepVariants)
head(df0)
```

```
##            mutation      PPI1      PPI2      PPI3    avePPI  logFC_shrunk
## 1 FOS.0.WT_JUN.0.WT 1.0000000 1.0000000 1.0000000 1.0000000  4.905382e-17
## 2  FOS.0.WT_JUN.1.* 0.4210242 0.4555306 0.3913326 0.4226291 -6.666359e+00
## 3  FOS.0.WT_JUN.1.A 0.9441778 0.9366920 0.9345138 0.9384612 -7.557261e-01
## 4  FOS.0.WT_JUN.1.C 0.9300885 0.9145116 0.9234750 0.9226917 -9.504132e-01
## 5  FOS.0.WT_JUN.1.D 0.9383403 0.9336526 0.9426828 0.9382252 -7.588861e-01
## 6  FOS.0.WT_JUN.1.E 0.9712508 0.9773245 0.9592317 0.9692690 -3.756026e-01
##      logCPM           FDR keepVariants
## 1 13.060466  1.000000e+00         TRUE
## 2  3.107818 7.802829e-208        FALSE
## 3  7.454886 4.129960e-132         TRUE
## 4  5.682072  1.272439e-57         TRUE
## 5  5.915706  1.899240e-43         TRUE
## 6  6.709633  2.071714e-19         TRUE
```

#### 9.1 Distribution of PPI scores for the individual replicates

The figure below shows the distribution of PPI scores across the variants retained in the paper, for each of the three replicates.

```
(g6 <- ggplot(df0 %>% dplyr::filter(keepVariants) %>%
                  dplyr::select(mutation, PPI1, PPI2, PPI3) %>%
                  tidyr::gather(key = Replicate, value = PPI, -mutation),
              aes(x = PPI, color = Replicate)) + 
     geom_line(stat = "density", size = 1) + theme_bw() +
     labs(x = "PPI score") + 
     theme(axis.text = element_text(size = 15),
           axis.title = element_text(size = 15)))
```

#### 9.2 Correlation between PPI scores for two replicates

Next, we plot the agreement of the PPI scores between the first two replicates, distinguishing between variants with a mutation in only one of the proteins, and variants with mutations in both.

```
df <- df0 %>% 
    dplyr::filter(mutation != "FOS.0.WT_JUN.0.WT") %>%
    dplyr::mutate(mutationType = 
                      c("double", "single")[grepl(".0.", mutation, fixed = TRUE) + 1]) %>%
    dplyr::mutate(mutFos = c("", "Fos")[as.numeric(!grepl("FOS.0.", mutation, 
                                                          fixed = TRUE)) + 1],
                  mutJun = c("", "Jun")[as.numeric(!grepl("JUN.0.", mutation, 
                                                          fixed = TRUE)) + 1]) %>%
    tidyr::unite("mutationTypeFull", mutationType, mutFos, mutJun, 
                 sep = "", remove = FALSE)
df %>% dplyr::filter(keepVariants) %>% dplyr::group_by(mutationType) %>%
    dplyr::summarize(correlation = cor(PPI1, PPI2, use = "pairwise.complete.obs"))
```

```
## # A tibble: 2 × 2
##   mutationType correlation
##   <chr>              <dbl>
## 1 double             0.951
## 2 single             0.995
```

```
(g7 <- ggplot() + 
        geom_point(data = df %>% dplyr::filter(keepVariants & mutationType == "double"), 
                   aes(x = PPI1, y = PPI2, color = mutationType), alpha = 0.01, 
                   size = 0.5) +
        geom_point(data = df %>% dplyr::filter(keepVariants & mutationType == "single"), 
                   aes(x = PPI1, y = PPI2, color = mutationType), alpha = 1,
                   size = 0.5) + 
        geom_abline(slope = 1, intercept = 0) + theme_bw() + 
        scale_color_manual(values = c(double = "blue", single = "orange"), 
                           name = "") + 
        labs(x = "PPI score, replicate 1", y = "PPI score, replicate 2") + 
        theme(axis.text = element_text(size = 15),
              axis.title = element_text(size = 15)))
```

#### 9.3 Heatmap of single mutant PPI scores

Finally, we plot a heatmap of the average PPI scores for the single mutant, stratified by the heptad position.

```
## Get single mutations only
ppis_single <- ppis[grep(".0.", rownames(ppis), fixed = TRUE), ]
ppis_single <- ppis_single[rownames(ppis_single) != "FOS.0.WT_JUN.0.WT", ]
dim(ppis_single)
```

```
## [1] 1280    3
```

```
head(ppis_single)
```

```
##                  OUT_vs_IN_replR1 OUT_vs_IN_replR2 OUT_vs_IN_replR3
## FOS.0.WT_JUN.1.*        0.4210242        0.4555306        0.3913326
## FOS.0.WT_JUN.1.A        0.9441778        0.9366920        0.9345138
## FOS.0.WT_JUN.1.C        0.9300885        0.9145116        0.9234750
## FOS.0.WT_JUN.1.D        0.9383403        0.9336526        0.9426828
## FOS.0.WT_JUN.1.E        0.9712508        0.9773245        0.9592317
## FOS.0.WT_JUN.1.F        0.9346549        0.9333547        0.9229442
```

```
hm <- 
    data.frame(mutation = rownames(ppis_single), avePPI = rowMeans(ppis_single), 
               stringsAsFactors = FALSE) %>%
    dplyr::filter(mutation %in% keepVariants) %>%
    dplyr::mutate(mutation = gsub("FOS.0.WT_", "", mutation),
                  mutation = gsub("_JUN.0.WT", "", mutation)) %>%
    dplyr::mutate(protein = sapply(strsplit(mutation, "\\."), .subset, 1),
                  position = sapply(strsplit(mutation, "\\."), .subset, 2),
                  aminoacid = sapply(strsplit(mutation, "\\."), .subset, 3)) %>%
    dplyr::select(-mutation) %>%
    dplyr::mutate(position = as.factor(as.numeric(position))) %>%
    dplyr::filter(aminoacid != "*") %>%
    dplyr::mutate(aminoacid = factor(
        aminoacid, levels = rev(c("A", "L", "I", "V", "F", "W", "Y", "H", 
                                  "S", "T", "Q", "N", "D", "E", "K", "R", 
                                  "M", "C", "G", "P"))))
(g8 <- ggplot(hm,
              aes(x = position, y = aminoacid, fill = avePPI)) + 
        geom_tile() + facet_wrap(~protein, ncol = 1) + 
        theme_bw() + 
        scale_fill_gradientn(colours = c("blue", "white", "darkred"), 
                             values = rescale(c(0.4, 1, 1.1)),
                             guide = "colorbar", limits = c(0.4, 1.1), 
                             na.value = "white", 
                             name = "Average\nPPI") + 
        geom_vline(xintercept = c(7.5, 14.5, 21.5, 28.5)) + 
        ylab("Amino acid") + 
        theme(panel.grid.major = element_blank(), panel.grid.minor = element_blank(),
              axis.title = element_text(size = 16), 
              strip.text = element_text(size = 14)))
```

### 10 Generate summary figure for paper

```
a1 <- cowplot::plot_grid(g1, g2, ncol = 1, align = "v", axis = "lr", 
                         rel_heights = c(1, 1), labels = c("A", "B"))
```

```
## Warning: Groups with fewer than two data points have been dropped.
```

```
a2 <- cowplot::plot_grid(g3 + theme(axis.text.x = element_text(size = 8), 
                                    axis.text.y = element_text(size = 8)),
                         g4 + theme(axis.text.x = element_text(size = 8), 
                                    axis.text.y = element_text(size = 8)),
                         nrow = 1, align = "h", axis = "b", rel_widths = c(1, 1), 
                         labels = c("C", "D"))
a3 <- cowplot::plot_grid(a1, a2, rel_widths = c(0.5, 1))
a4 <- cowplot::plot_grid(ggmatrix_gtable(g5), g8, nrow = 1, align = "h", 
                         axis = "b", rel_widths = c(1, 1), labels = c("E", "F"))
cowplot::plot_grid(a3, a4, ncol = 1)
```

### 11 Session info

```
sessionInfo()
```

```
## R version 4.2.1 (2022-06-23)
## Platform: x86_64-pc-linux-gnu (64-bit)
## Running under: CentOS Linux 7 (Core)
## 
## Matrix products: default
## BLAS/LAPACK: /tungstenfs/groups/gbioinfo/Appz/easybuild/software/OpenBLAS/0.3.12-GCC-10.2.0/lib/libopenblas_skylakex-r0.3.12.so
## 
## locale:
##  [1] LC_CTYPE=en_US.UTF-8       LC_NUMERIC=C              
##  [3] LC_TIME=en_US.UTF-8        LC_COLLATE=en_US.UTF-8    
##  [5] LC_MONETARY=en_US.UTF-8    LC_MESSAGES=en_US.UTF-8   
##  [7] LC_PAPER=en_US.UTF-8       LC_NAME=C                 
##  [9] LC_ADDRESS=C               LC_TELEPHONE=C            
## [11] LC_MEASUREMENT=en_US.UTF-8 LC_IDENTIFICATION=C       
## 
## attached base packages:
## [1] stats4    stats     graphics  grDevices utils     datasets  methods  
## [8] base     
## 
## other attached packages:
##  [1] GGally_2.1.2                cowplot_1.1.1              
##  [3] ggplot2_3.3.6               scales_1.2.1               
##  [5] SummarizedExperiment_1.26.1 Biobase_2.56.0             
##  [7] GenomicRanges_1.48.0        GenomeInfoDb_1.32.4        
##  [9] IRanges_2.30.1              S4Vectors_0.34.0           
## [11] BiocGenerics_0.42.0         MatrixGenerics_1.8.1       
## [13] matrixStats_0.62.0          mutscan_0.2.31             
## 
## loaded via a namespace (and not attached):
##  [1] sass_0.4.2             edgeR_3.38.4           tidyr_1.2.1           
##  [4] splines_4.2.1          jsonlite_1.8.0         bslib_0.4.0           
##  [7] assertthat_0.2.1       highr_0.9              GenomeInfoDbData_1.2.8
## [10] Rsamtools_2.12.0       yaml_2.3.5             ggrepel_0.9.1         
## [13] pillar_1.8.1           lattice_0.20-45        glue_1.6.2            
## [16] limma_3.52.3           digest_0.6.29          RColorBrewer_1.1-3    
## [19] XVector_0.36.0         colorspace_2.0-3       htmltools_0.5.3       
## [22] Matrix_1.5-1           plyr_1.8.7             pkgconfig_2.0.3       
## [25] csaw_1.30.1            bookdown_0.29          zlibbioc_1.42.0       
## [28] purrr_0.3.4            BiocParallel_1.30.3    tibble_3.1.8          
## [31] farver_2.1.1           generics_0.1.3         grr_0.9.5             
## [34] ellipsis_0.3.2         DT_0.25                cachem_1.0.6          
## [37] withr_2.5.0            cli_3.4.1              magrittr_2.0.3        
## [40] crayon_1.5.1           evaluate_0.16          fansi_1.0.3           
## [43] tools_4.2.1            lifecycle_1.0.2        stringr_1.4.1         
## [46] munsell_0.5.0          locfit_1.5-9.6         DelayedArray_0.22.0   
## [49] Biostrings_2.64.1      compiler_4.2.1         jquerylib_0.1.4       
## [52] rlang_1.0.6            grid_4.2.1             Matrix.utils_0.9.8    
## [55] RCurl_1.98-1.8         htmlwidgets_1.5.4      labeling_0.4.2        
## [58] bitops_1.0-7           rmarkdown_2.16         gtable_0.3.1          
## [61] codetools_0.2-18       DBI_1.1.3              reshape_0.8.9         
## [64] R6_2.5.1               knitr_1.40             dplyr_1.0.10          
## [67] fastmap_1.1.0          utf8_1.2.2             metapod_1.4.0         
## [70] stringi_1.7.8          parallel_4.2.1         Rcpp_1.0.9            
## [73] vctrs_0.4.1            tidyselect_1.1.2       xfun_0.33
```
