## Additional file 2 for "mutscan - a flexible R package for efficient end-to-end analysis of multiplexed assays of variant effect data"

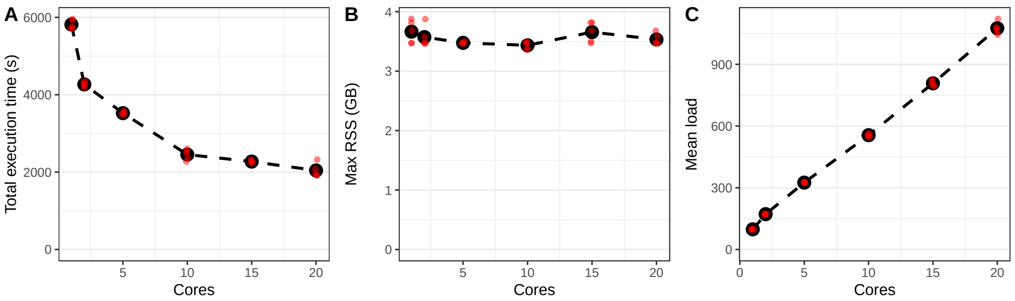


Figure S1: Computational performance metrics for *mutscan*'s *digestFastqs()* function run with different numbers of cores, processing a single input sample from the Li_tRNA_sel30 dataset. The black dots represent the average across five independent runs, each indicated by a smaller red dot. The dashed curves connect the average values for different numbers of cores. RSS - resident set size.

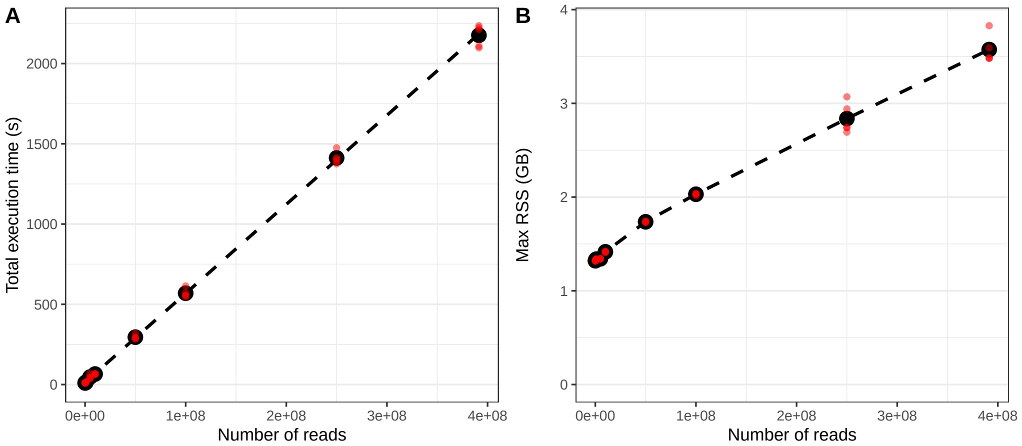


Figure S2: Execution time and maximum memory required by the *digestFastqs()* function when processing different numbers of reads (achieved by setting the `maxNReads` argument of the *digestFastqs()* function to `N`, which limits the processing to the first `N` reads in the FASTQ file). The black dots represent the average across five independent runs, each indicated by a smaller red dot. The dashed line in (A) is a linear regression line, while the dashed curve in (B) connects the average values for different numbers of cores. RSS - resident set size.

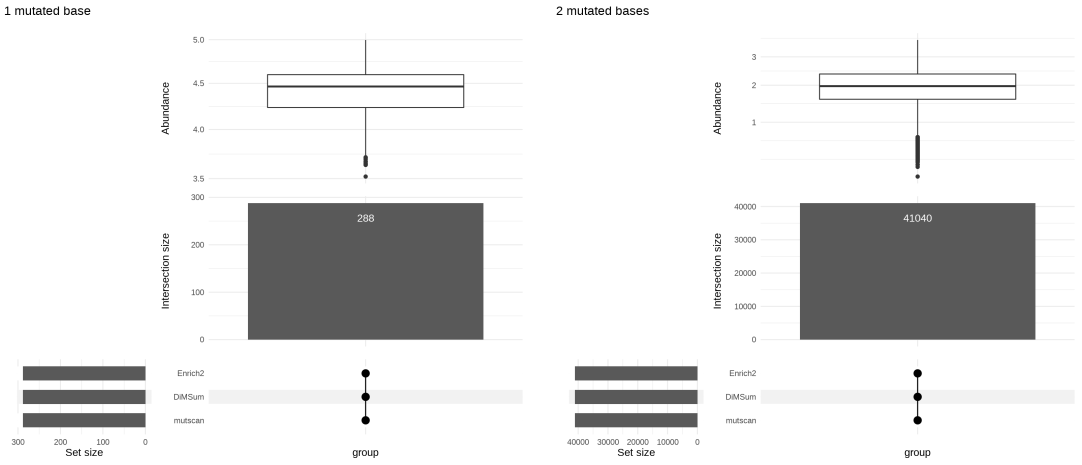


Figure S3: Comparison of the variants detected by *mutscan*, *DiMSum* and *Enrich2* in the Diss_FOS dataset, stratified by the number of mutated bases in the variant. All variants with up to two mutated bases are consistently detected by all three tools. The abundance represents the average $log_{10}(count + 1)$ across samples where the variant was quantified.

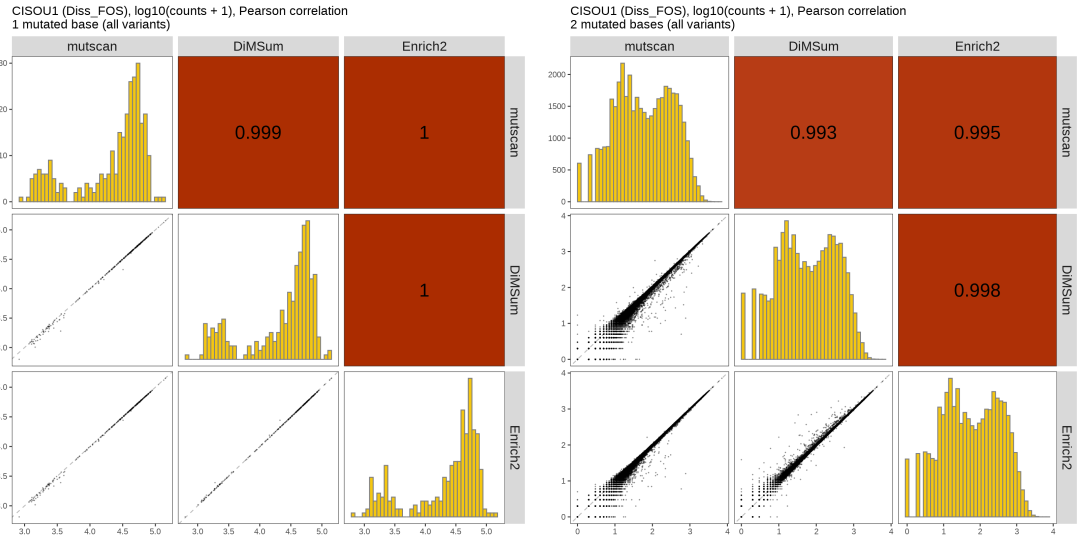


Figure S4: Comparison of the observed counts for variants detected by *mutscan*, *DiMSum* and *Enrich2* in the Diss_FOS dataset, stratified by the number of mutated bases in the variant.

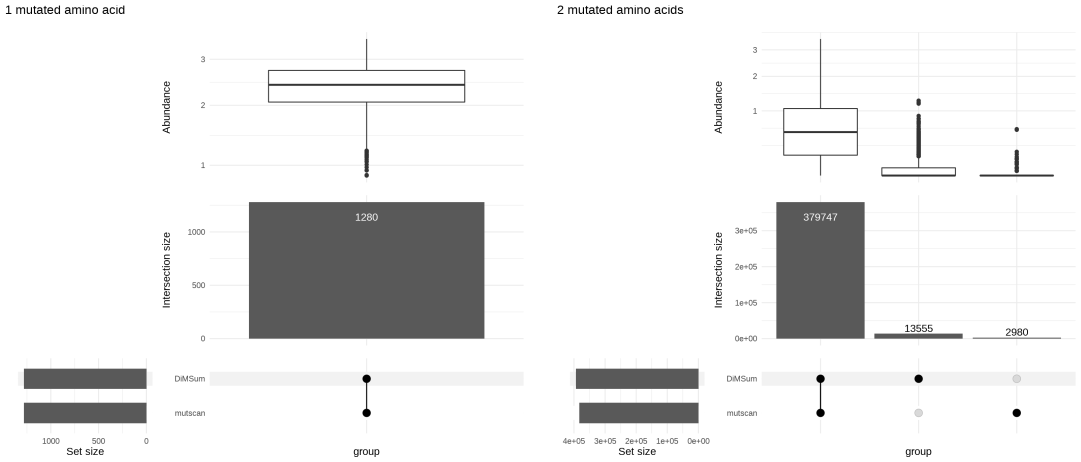


Figure S5: Comparison of the variants detected by *mutscan* and *DiMSum* in the Diss_FOS_JUN dataset, stratified by the number of mutated amino acids in the variant. Most variants are found consistently with both tools. The ones found by a single tool tend to have a low read count. The abundance represents the average $log_{10}(count + 1)$ across samples where the variant was quantified.

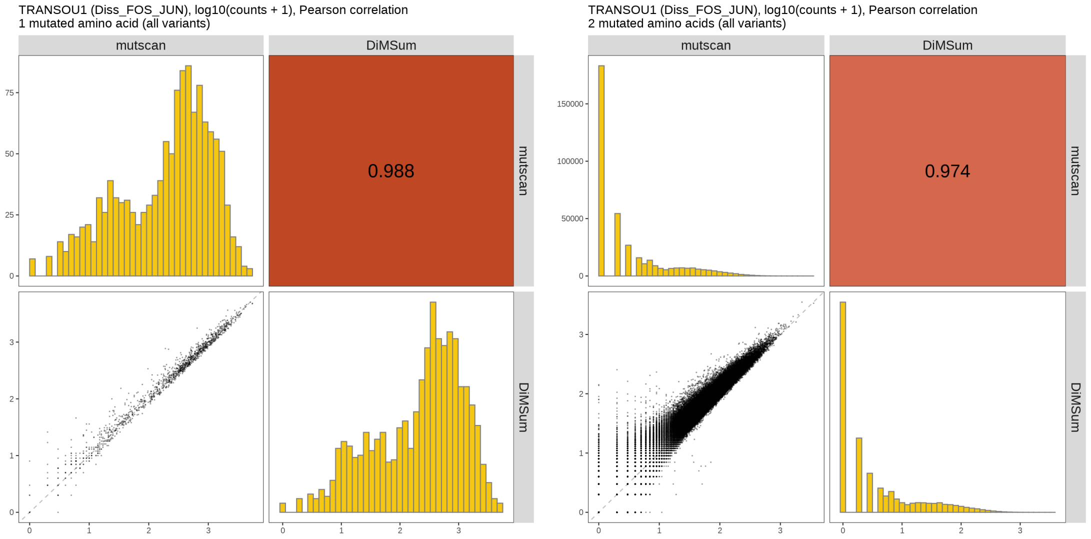


Figure S6: Comparison of the observed counts for variants detected by *mutscan* and *DiMSum* in the Diss_FOS_JUN dataset, stratified by the number of mutated amino acids in the variant.

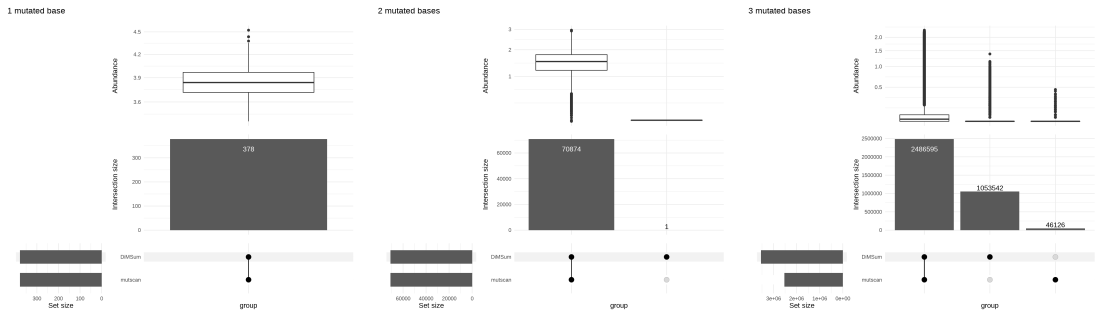

Figure S7: Comparison of the variants detected by *mutscan* and *DiMSum* in the Bolognesi_TDP43_290_331 dataset, stratified by the number of mutated bases in the variant. Almost all variants with up to two mutations are consistently detected by both methods. The ones found by a single tool tend to have a low read count. The abundance represents the average $log_{10}(count + 1)$ across samples where the variant was quantified.

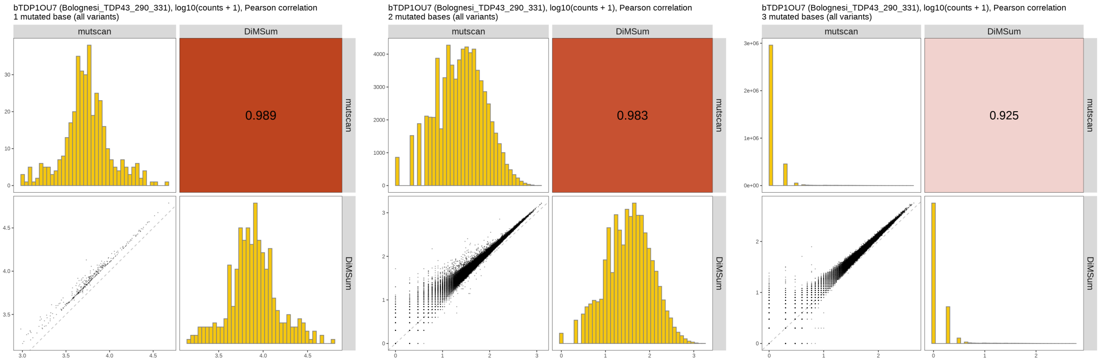

Figure S8: Comparison of the observed counts for variants detected by *mutscan* and *DiMSum* in the Bolognesi_TDP43_290_331 dataset, stratified by the number of mutated bases in the variant.

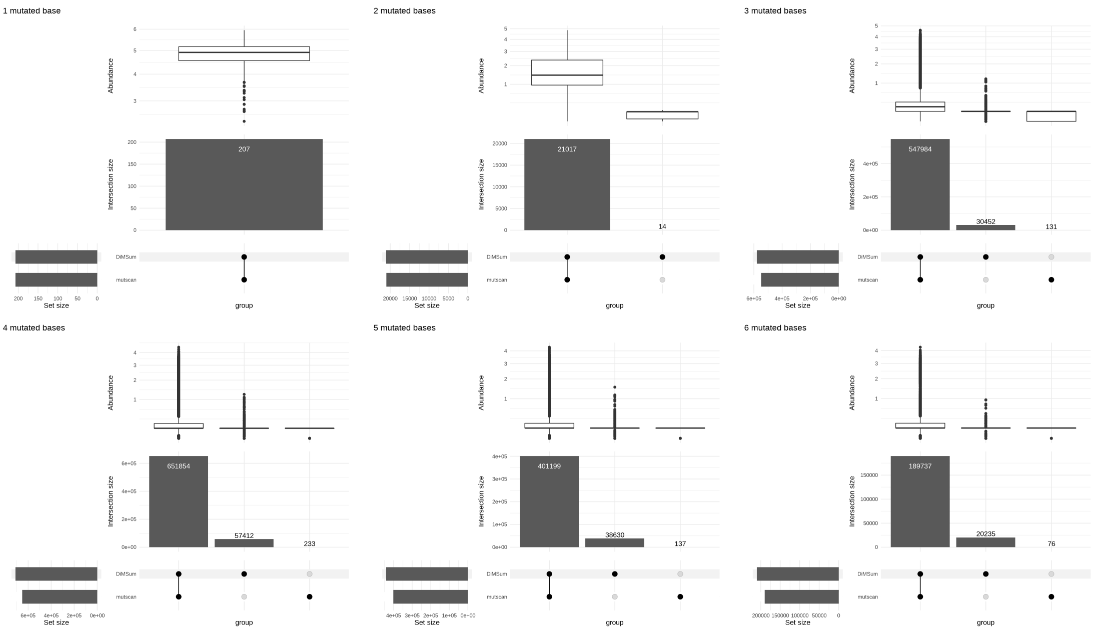


Figure S9: Comparison of the variants detected by *mutscan* and *DiMSum* in the Li_tRNA_sel30 dataset, stratified by the number of mutated bases in the variant. Only variants with up to six mutations are shown. Most variants are found with both tools, and the ones found by a single tool tend to have a lower read count. The abundance represents the average $log_{10}(count + 1)$ across samples where the variant was quantified.

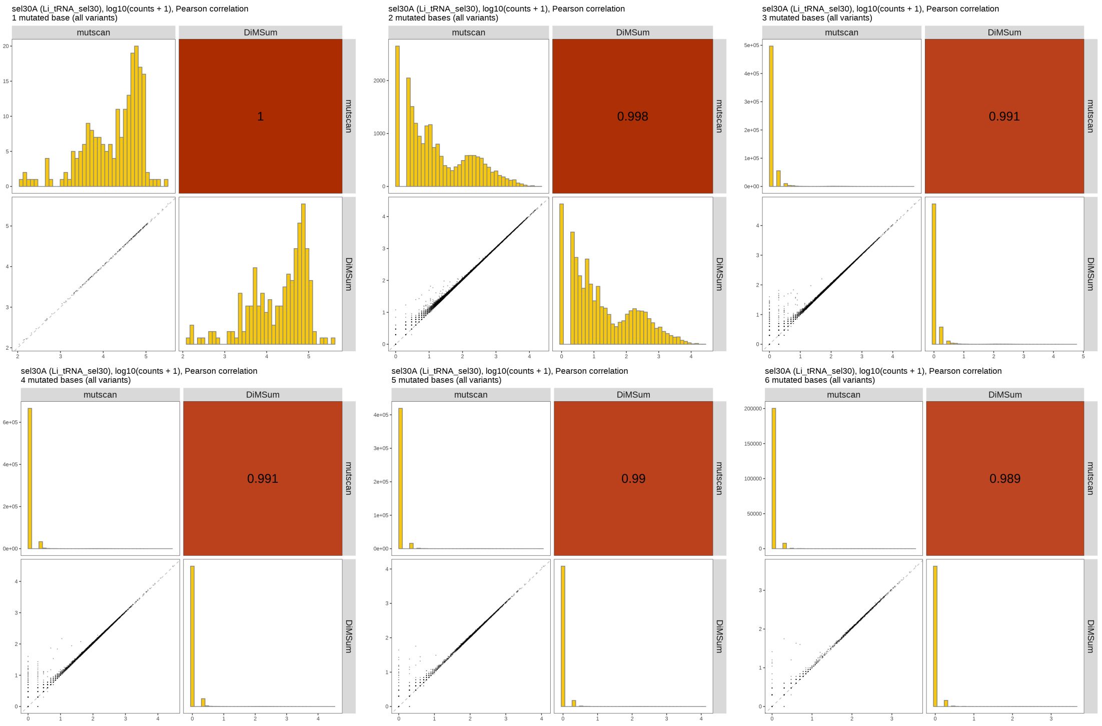


Figure S10: Comparison of the observed counts for variants detected by *mutscan* and *DiMSum* in the Li_tRNA_sel30 dataset, stratified by the number of mutated bases in the variant. Only variants with up to six mutations are shown.

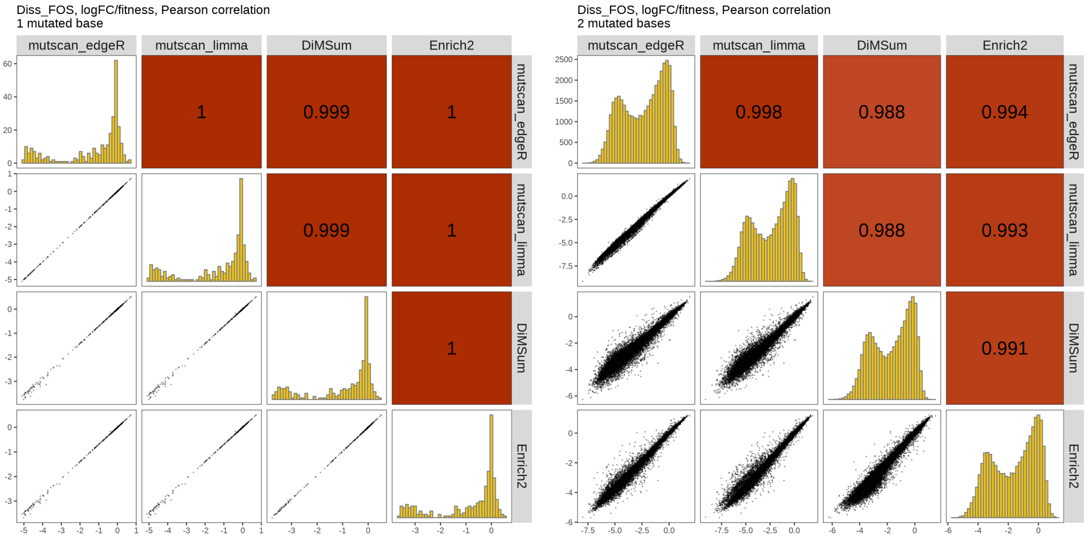


Figure S11: Comparison of fitness scores estimated by *mutscan*, *DiMSum* and *Enrich2* in the Diss_FOS dataset, stratified by the number of mutated bases in the variant. The agreement between the fitness scores from the different methods is very high for variants with a single mutation, and decreases as the number of mutations increases (and alongside that, the average abundance decreases).

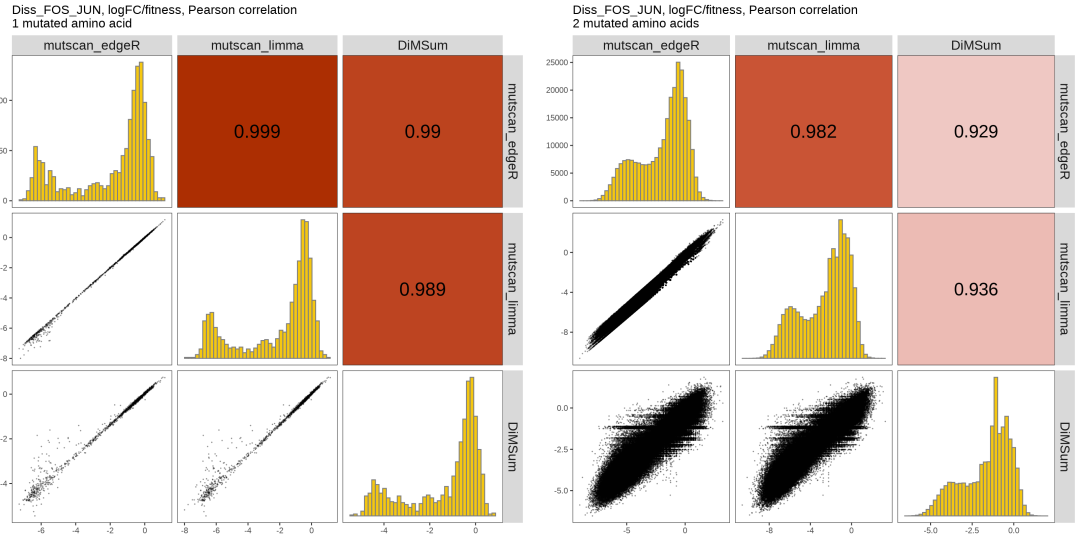


Figure S12: Comparison of fitness scores estimated by *mutscan* and *DiMSum* in the Diss_FOS_JUN dataset, stratified by the number of mutated amino acids in the variant. The agreement between the fitness scores from the different methods is very high for variants with a single mutation, and decreases as the number of mutations increases.

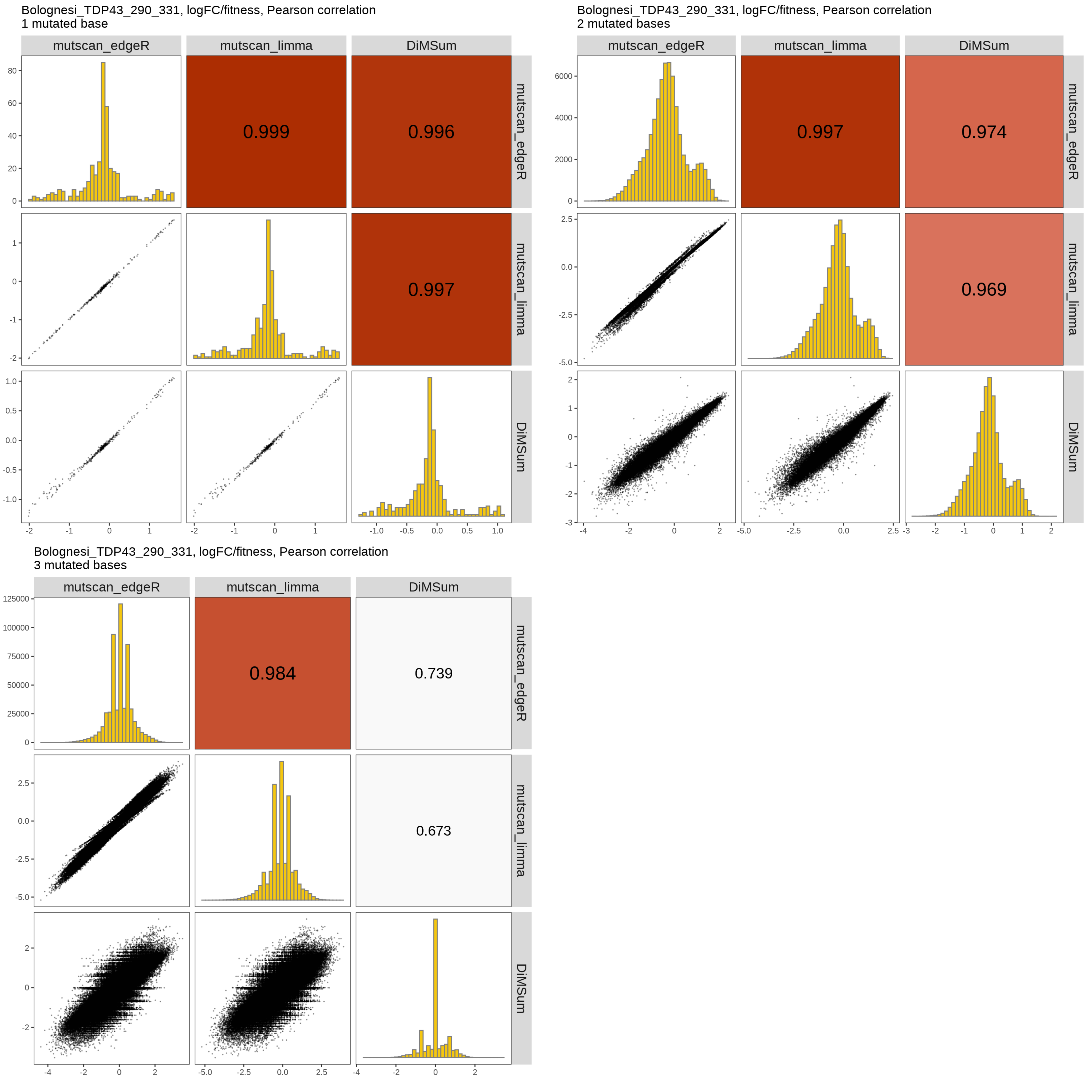


Figure S13: Comparison of fitness scores estimated by *mutscan* and *DiMSum* in the Bolognesi_TDP43_290_331 dataset, stratified by the number of mutated bases in the variant. The agreement between the fitness scores from the different methods is very high for variants with a single mutation, and decreases as the number of mutations increases.

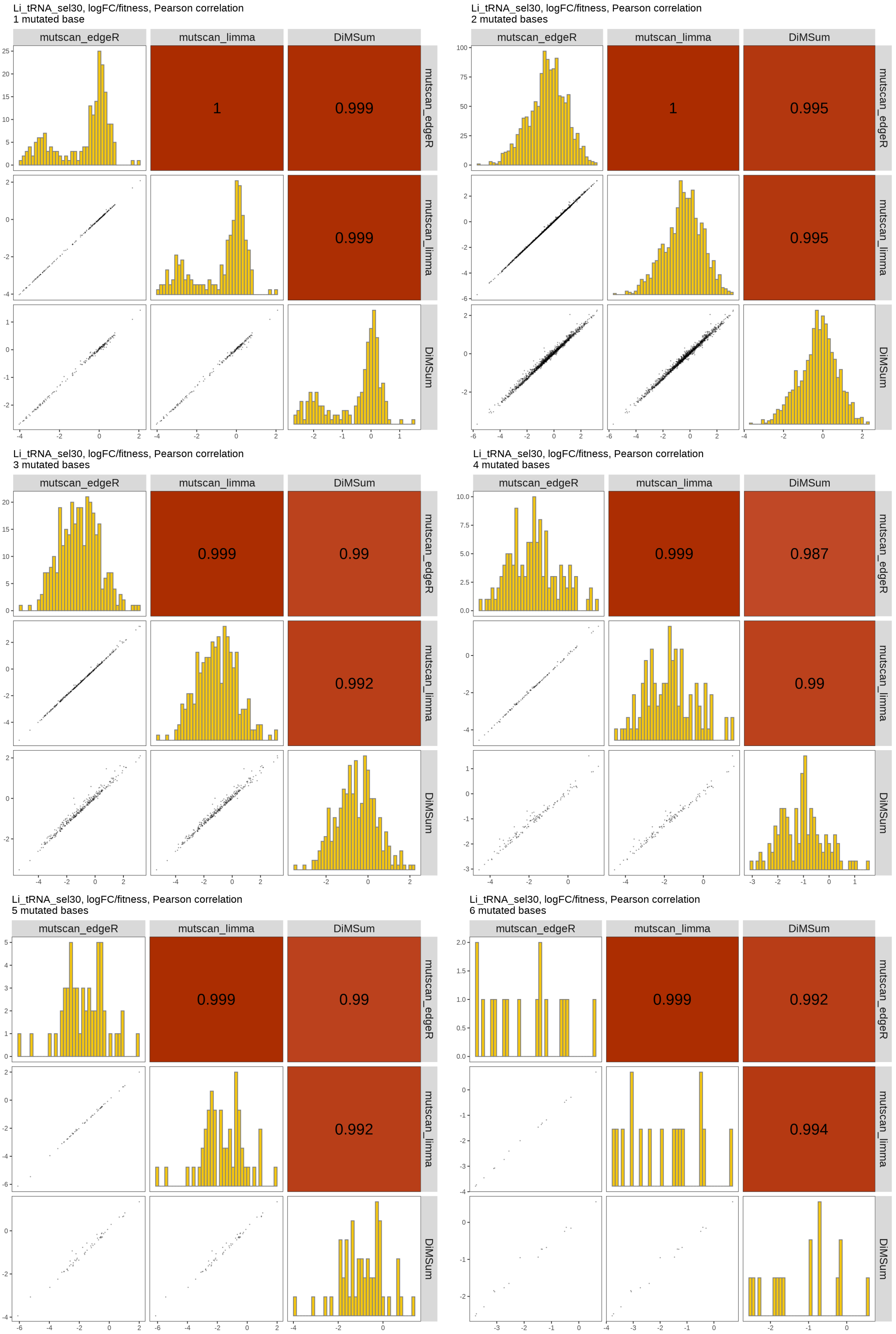


Figure S14: Comparison of fitness scores estimated by *mutscan* and *DiMSum* in the Li_tRNA_sel30 dataset, stratified by the number of mutated bases in the variant.

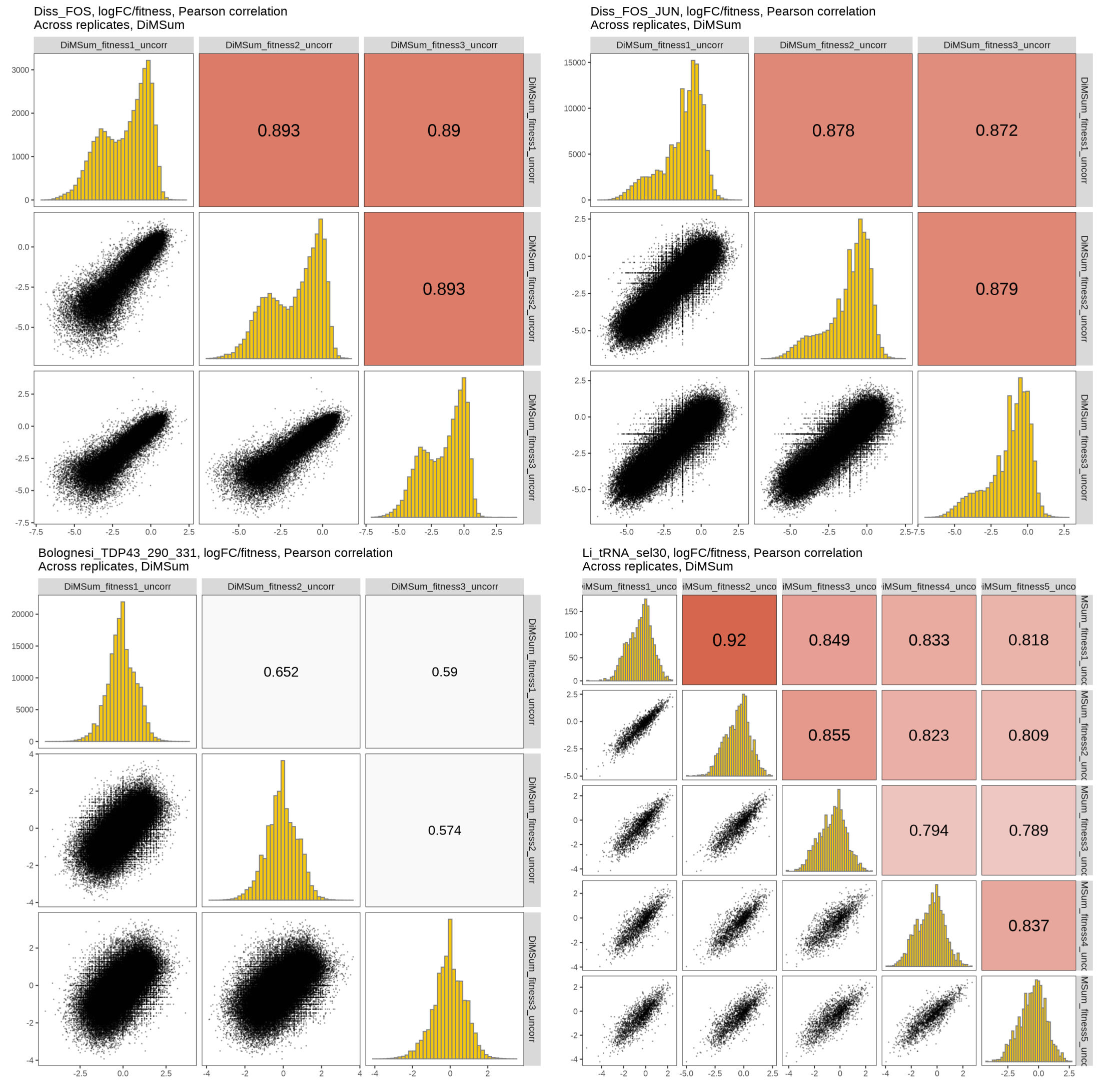


Figure S15: Comparison of fitness scores estimated by *DiMSum* for individual replicates in the four example data sets.
